# Human ACE2 receptor polymorphisms predict SARS-CoV-2 susceptibility

**DOI:** 10.1101/2020.04.07.024752

**Authors:** Eric W. Stawiski, Devan Diwanji, Kushal Suryamohan, Ravi Gupta, Frederic A. Fellouse, J. Fah Sathirapongsasuti, Jiang Liu, Ying-Ping Jiang, Aakrosh Ratan, Monika Mis, Devi Santhosh, Sneha Somasekar, Sangeetha Mohan, Sameer Phalke, Boney Kuriakose, Aju Antony, Jagath R. Junutula, Stephan C. Schuster, Natalia Jura, Somasekar Seshagiri

**Author notes:** correspondence: NJ - and SS –. co-first author (Eric W. Stawiski, Devan Diwanji, Kushal Suryamohan).

## Abstract

Severe acute respiratory syndrome coronavirus 2 (SARS-CoV-2) is the cause of coronavirus disease (COVID-19) that has resulted in a global pandemic. It is a highly contagious positive strand RNA virus and its clinical presentation includes severe to critical respiratory disease that appears to be fatal in ∼3-5% of the cases. The viral spike (S) coat protein engages the human angiotensin-converting enzyme2 (ACE2) cell surface protein to invade the host cell. The SARS-CoV-2 S-protein has acquired mutations that increase its affinity to human ACE2 by ∼10-15-fold compared to SARS-CoV S-protein, making it highly infectious. In this study, we assessed if ACE2 polymorphisms might alter host susceptibility to SARS-CoV-2 by affecting the ACE2 S-protein interaction. Our comprehensive analysis of several large genomic datasets that included over 290,000 samples representing >400 population groups identified multiple ACE2 protein-altering variants, some of which mapped to the S-protein-interacting ACE2 surface. Using recently reported structural data and a recent S-protein-interacting synthetic mutant map of ACE2, we have identified natural ACE2 variants that are predicted to alter the virus-host interaction and thereby potentially alter host susceptibility. In particular, human ACE2 variants S19P, I21V, E23K, K26R, T27A, N64K, T92I, Q102P and H378R are predicted to increase susceptibility. The T92I variant, part of a consensus NxS/T N-glycosylation motif, confirmed the role of N90 glycosylation in immunity from non-human CoVs. Other ACE2 variants K31R, N33I, H34R, E35K, E37K, D38V, Y50F, N51S, M62V, K68E, F72V, Y83H, G326E, G352V, D355N, Q388L and D509Y are putative protective variants predicted to show decreased binding to SARS-CoV-2 S-protein. Overall, ACE2 variants are rare, consistent with the lack of selection pressure given the recent history of SARS-CoV epidemics, however, are likely to play an important role in altering susceptibility to CoVs.

## Introduction

Coronaviruses (CoVs) are widely distributed in nature and pose a serious threat to humans and a range of mammalian hosts causing respiratory, gastrointestinal, and central nervous system diseases (Li, 2016). CoVs are enveloped non-segmented positive-sense single stranded RNA viruses and are classified into α−, β−, γ−, and δ-CoVs (Li, 2016). While α- and β-CoVs infect mammals, the γ- and δ-CoVs generally infect birds (Li, 2016). Previously, α-CoVs HCoV-229E and HCoV-NL63, and β-CoVs HCoV-HKU1 and HCoV-OC43 have been found to infect humans leading to mild symptoms (Graham and Baric, 2010; Li, 2016). However, three β-CoVs, severe acute respiratory syndrome coronavirus (SARS-CoV) in 2003 (Holmes, 2003; Li, 2016), Middle-East respiratory syndrome coronavirus in 2012 (MERS-CoV) (Li, 2016; Zaki et al., 2012), and more recently SARS-CoV-2 in 2019 (Chan et al., 2020; Huang et al., 2020; Zhu et al., 2020), have crossed species barrier to infect humans resulting in respiratory illnesses including pneumonia that is fatal.

SARS-CoV-2 is a novel coronavirus (2019-nCoV) first reported in December 2019 and is the cause of an ongoing global pandemic (Chan et al., 2020; Huang et al., 2020; Zhu et al., 2020). It has infected over 1.2 million people in 181 countries leading to over 69,000 deaths as of April 5^th^, 2020 (JHU, 2020). SARS-CoV-2 genome sequence analysis revealed that it is closer to bat CoV RaTG13 (96.2% identical) than to SARS-CoV (79.5% identical) responsible for the 2003 epidemic, suggesting that this novel virus originated in bats independently before jumping to humans either directly or through an yet to be determined intermediary host (Guo et al., 2020).

As with SARS-CoV and a related alpha-coronavirus NL63 (HCoV-NL63), SARS-CoV-2 employs the human ACE2 cell surface protein as a receptor to gain entry into cells (Hoffmann et al., 2020; Letko et al., 2020; Lin et al., 2008; Ou et al., 2020; Wan et al., 2020; Zhou et al., 2020a). The virus surface spike glycoprotein (S-protein) constitutes a key determinant of viral host range and contains two domains, S1 and S2, which are separated by a protease cleavage site (Li, 2016). A successful host cell invasion by the virus involves direct binding of the virus S1 receptor binding domain (RBD) to the host ACE2 peptidase extracellular domain (PD), exposing the S1-S2 inter-domain protease site that upon cleavage by host proteases leads to S2-mediated virus-host cell membrane fusion (Belouzard et al., 2009; Hoffmann et al., 2020; Li, 2016; Li et al., 2005a; Simmons et al., 2005).

The SARS-CoV-2 S-protein is 98% identical to the bat CoV RaTG13 S protein, with the exception of an insertion that is also absent in the SARS-CoV S-protein in the S1/S2 inter-domain protease cleavage site. This difference has been proposed to alter SARS-CoV-2 tropism and enhance its transmissibility (Walls et al., 2020).

Several structural studies involving the SARS-CoV-2 S-protein RBD and ACE2 PD have identified key residues involved in their interaction (Shang et al., 2020; Walls et al., 2020; Wrapp et al., 2020; Yan et al., 2020). The S-protein RBD was reported to bind ACE2 PD with ∼10- to 20-fold higher affinity (∼15 nM) when compared to the SARS-CoV S-protein RBD (Shang et al., 2020; Wrapp et al., 2020), potentially contributing to high rate of SARS-CoV-2 infection.

As the interactions between the ACE2 receptor and S-protein RBD interface are critical for the cellular entry of the virus, we wondered if there were natural ACE2 variations that decrease or increase its affinity to the S-protein RBD that may protect or render individuals more susceptible to the virus. Consistent with this possibility, a saturation mutagenesis screen of select ACE2 PD residues identified variants that showed enhanced or decreased binding to S-protein (Procko, 2020).

In this study, we have analyzed ACE2 protein-altering variants in a large cohort of human population groups and identified polymorphisms that either likely protect or render individuals more susceptible to the virus. Understanding these changes at the molecular level, combined with the genotype and epidemiological data will allow the elucidation of population risk profiles and also help advance therapeutics such as a rationally designed soluble ACE2 receptor for treatment of COVID-19.

## Results

### Human ACE2 population polymorphism

SARS-CoV-2 S-protein interacts with the ACE2 PD to enter the human host cells. Analysis of S-protein RBD domain of SARS-CoV-2, SARS-CoV and closely related bat CoV RaTG13 identified changes that have increased the affinity of CoV-2 S1 RBD for human ACE2, which likely contributes to its increased infectivity (Shang et al., 2020; Wrapp et al., 2020). It is very likely that there are natural variations in ACE2 in human populations, though not under selection, that may increase or decrease its affinity to SARS-CoV-2 S-protein and thereby render individuals more resistant or susceptible to the virus. To investigate this, we assessed ACE2 protein-altering variations from a number of databases including the gnomAD (Karczewski et al., 2019), RotterdamStudy (Ikram et al., 2017), ALSPAC (Fraser et al., 2013) and Asian-specific databases which included GenomeAsia100k (GenomeAsia, 2019), HGDP (Bergstrom et al., 2020), TOMMO-3.5kjpnv2 (Tadaka et al., 2019), and IndiGen (https://indigen.igib.in/), and HGDP (Bergstrom et al., 2020) (Supplementary Table 1). We found a total of 298 unique protein altering variants across 256 codons distributed throughout the 805 amino acid long human ACE2 (**Figure 1a, 1b, Supplementary Figure 1**, and **Supplementary Table 1**). The most frequent variant, N720D (1.6% allele frequency; n=3054, gnomAD), was found in the C-terminal collectrin domain that is not involved in the SARS-CoV-2 S-protein interaction. Overall, we found human ACE2 receptor polymorphisms to be low with a weighted mean Fst (fixation index) value of 0.0168, and the ACE2 PD showed even more reduced variation (Wilcoxon p=0.0656, **Supplementary Figure 2a**, see Methods). Further, we found ACE2 to be highly intolerant of loss of function variants (pLI=0.9977, gnomAD; **Supplementary Figure 2b**, see Methods), though we observed 5 predicted LOF singleton alleles (**Supplementary Table 1**).

**Figure 1.**
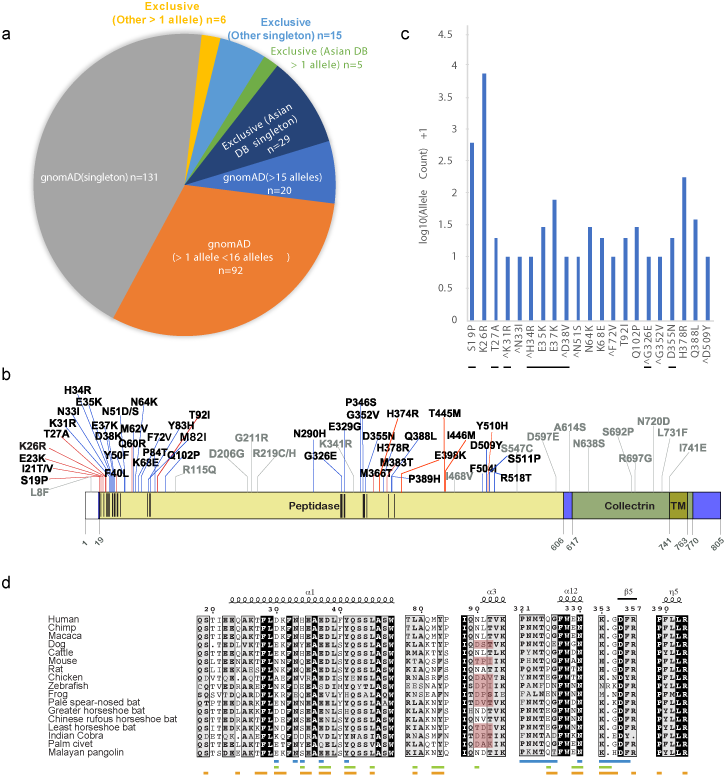
ACE2 polymorphisms. **a**. Pie chart representing protein altering variations in ACE2 by allele count and source. **b**. ACE2 protein domain showing positions with polymorphisms that were predicted to lead to increased (red line) or decreased (blue line) S-protein binding. Recurrent polymorphisms (n>1) that were predicted to not impact S-protein binding are show in light grey. Residues within the ACE2 PD known to interact with viral S-protein are shown as black vertical lines within the peptidase domain in the ACE2 diagram. **c**. Log base 10 pseudo count adjusted (+1) observed ACE2 allele counts of mutants predicted to impact S-protein binding. Singletons are marked with a ^ and direct S-protein contact residues are underlined. **d**. Multiple sequence alignment of the S-protein interacting ACE2 sequence from indicated species. ACE2 NxT/S glycosylation motif disrupted in dog, rat, palm civet and several bat ACE2 is highlighted in red. ACE2 residues that mediate contact with NL63-CoV, SARS-CoV and SARS-CoV-2 are shown as blue, green and orange bars, respectively.

**Figure 2.**
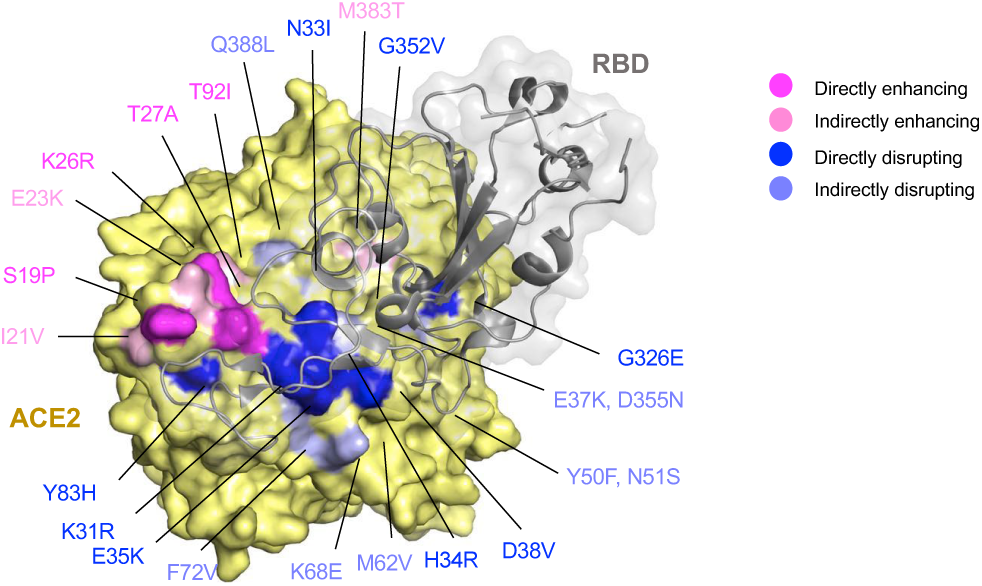
Identified polymorphisms in human ACE2 mapped to the structure of human ACE2 in complex with the SARS-CoV-2 RBD. Residues in ACE2 showing polymorphic variation in human population were mapped on to the structure of the ACE2/SARS-CoV-2 RBD (PDB: 6VW1) and colored according to their effect on the predicted affinity between human ACE2 and viral spike S-protein systematically as measured by a deep sequence mutagenesis screen (Procko, 2020). Polymorphisms that were predicted to enhance the binding between ACE2 and the S-protein are colored in magenta if they are located directly at the ACE2/SARS-CoV-2 interface or in light pink if they do not participate directly in the interface. Polymorphisms that are predicted to enhance the binding between ACE2 and the S-protein are colored in dark blue if they are located directly at the ACE2/SARS-CoV-2 interface or in light blue if they do not participate directly in the interface interactions.

Structural studies involving SARS-CoV and, more recently, the SARS-CoV-2 S-protein and its complex with human ACE2 have identified three regions in an ∼120 amino acid claw-like exposed outer surface of the human ACE2 (ACE2-claw) that contributes to its binding to the S-protein (Shang et al., 2020; Walls et al., 2020; Wrapp et al., 2020; Yan et al., 2020). The main residues at the interface include S19, Q24, T27, F28, D30, K31, H34, E35, E37, D38, Y41, Q42, L45, L79, M82, Y83, T324, Q325, G326, E329, N330, K353, G354, D355, R357, P389, and R393 (**Figure 1b**). Mutagenesis of four residues in the S-protein-binding interface of rat ACE2 was sufficient to convert rat ACE2 into a human SARS-CoV receptor, further indicating the importance of this region in determining the host range and specificity of CoVs (Li et al., 2005b). Considering these findings, we focused on variants within the human ACE2-claw S-protein RBD-binding interface and identified protein alterations in 44 codons that resulted in 49 unique variants for a total of 968 allelic variants. This included K26R, the second most frequent human ACE2 protein-altering variant (0.4% allele frequency; allele count=797, gnomAD), S19P, T27A, K31R, N33I, H34R, E35K, E37K, D38V, N51S, N64K, K68E, F72V, T921, Q102P, G326E, G352V, D355N, H378R, Q388L, and D509Y (**Supplementary Table 2**). These variants are likely to either increase or decrease the binding affinity of ACE2 to the S-protein and thereby alter the ability of the virus to infect the host cell.

A recent mutagenesis screen using a synthetic human ACE2 mutant library identified variants that either increased or decreased its binding to SARS-CoV-2 S-protein (Procko, 2020). Using a sequencing-based enrichment assay, the fold enrichment or depletion of the mutant sequences was measured in this study (Procko, 2020). Mapping the enrichment z-scores from this study (Procko, 2020) to the spectrum of natural ACE2 polymorphisms, we identified several rare ACE2 variants (**Figure 1c**) that likely alter their binding to the SARS-CoV-2 S-protein and thereby protect or render individuals more susceptible to the virus(Supplementary Table 2). The majority of the variants that were predicted to alter the interaction between ACE2 and the virus S-protein were clustered around the N-terminal region of ACE2 that interacts with the S-protein (**Figure 1b**).

Included among the ACE2 polymorphic variants that increase ACE2/S-protein interaction are S19P, I21V, E23K, K26R, K26E, T27A, N64K, T92I, Q102P, M383T and H378R (**Supplementary Table 2 and Supplementary Figure 3**). Among these, the T92I polymorphism stands out in particular because it is part of a NxT/S (where x is any amino acid except proline) consensus N-glycosylation motif (Gavel and von Heijne, 1990) where N90 is the site of N-glycan addition. The ACE2 NxT/S motif, while conserved in 96 out of 296 jawed vertebrate with ACE2 sequence available is absent or altered in several species, including the civet cat (*Paguma larvata*) and several bat species where residue N90 is mutated, a proline is present at position 91 or the T92 is altered to any amino acid except serine (**Figure 1d, Supplementary Figure 4 and Supplementary Table 3**) (Demogines et al., 2012; Gavel and von Heijne, 1990; Li et al., 2005b). These ACE2 variations are expected to abolish glycosylation at N90 (Gavel and von Heijne, 1990). Furthermore, a mutation that altered the NxT/S motif in human ACE2 to a civet ACE2-like sequence (90-NLTV-93 to DAKI), expected to abolish the N-glycosylation, increased the SARS-CoV infectivity and S-protein binding (**Figure 1d**) (Li et al., 2005b). The T92I mutant we identified showed a strong enrichment in the sequencing-based screen for S-protein binders (Procko, 2020). Considering these observations, we conclude that the T92I mutation increases the ACE2/S-protein binding affinity rendering individuals harboring this mutation more susceptibility to the virus.

**Figure 3.**
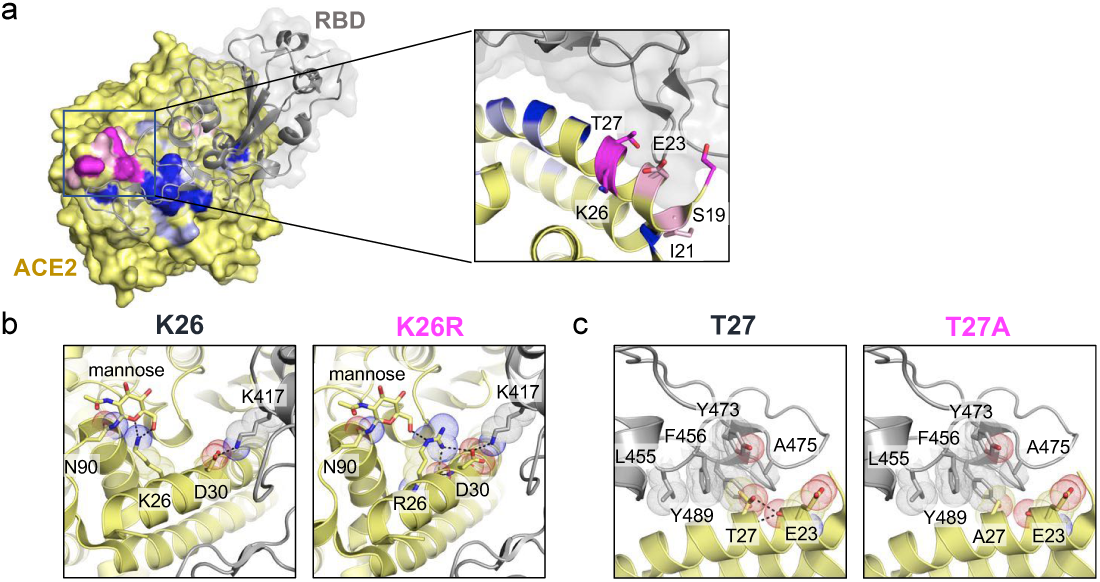
Structural basis of enhanced interaction between human ACE2 polymorphic variants and SARS-CoV-2 S-protein. **(a)** Zoomed-in view of the indicated polymorphic variants that enhance the interaction between the human ACE2 and the viral S-protein. The mutation color codes used are same as in Fig 1. **(b-c)** Analysis of the effect of the most frequent polymorphism, K26R (b) and T27A (c) polymorphisms. PDB structures shown are 6VW1 (a) and 6LZG (b,c).

**Figure 4.**
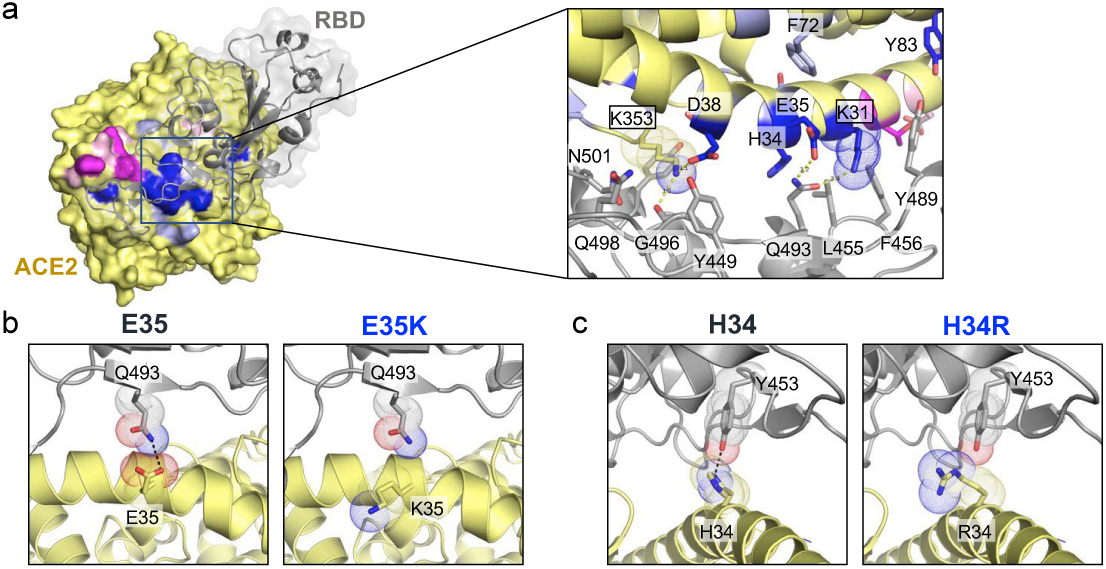
Structural basis of decreased interaction between human ACE2 polymorphic variants and SARS-CoV-2 S-protein. **(a)** Zoomed-in view of the indicated polymorphic variants that destabilize the interaction between human ACE2 and the viral S-protein. The mutation color codes used are same as in Fig 1.. Two hotspot lysine residues on ACE2 (K31 and K353) are highlighted with dot representation in the panel on the right. **(b)** Analysis of the recurring polymorphism at E35 of ACE2. **(c)** Analysis of the effect of ACE2 H34 mutation to arginine. PDB structures shown are 6VW1 (a) and 6LZG (b,c).

Variants that are predicted to reduce the virus S-protein interactions and thereby decrease S/ACE2 binding affinity include K31R, N33I, H34R, E35K, E37K, D38V, Y50F, N51S, K68E, F72V, Y83H, G326E, G352V, D355N and Q388L. Below we discuss the structural basis for the inhibitory effect on ACE2/S-protein binding for this selected set of mutations, as well as for the enhancing effect of the selected polymorphisms that were shown to increase ACE2/S-protein binding in vitro (Procko, 2020).

### Structural evaluation of ACE2 polymorphism

To further understand the effect of the polymorphisms on receptor recognition by the SARS-CoV-2 RBD and to confirm our predictions, we structurally modeled the identified ACE2 variants using the recently published cryo-EM and crystal structures of ACE2/SARS-CoV-2 RBD complexes (**Supplementary Table 4**) (Shang et al., 2020; Walls et al., 2020; Wrapp et al., 2020; Yan et al., 2020). We combined our structural analysis predictions with data from the Procko study (Procko, 2020) and classified the polymorphisms into five categories. These included mutations that directly enhanced or disrupted the residues at the ACE2/S-protein binding interface or residues interacting with the N90-linked glycan. We term these mutations as “directly enhancing” or “directly disrupting”. We also identified variants that were “indirectly enhancing” or “indirectly disrupting”. These mutants were found to affect the residues that are at the ACE2/S-protein binding interface but not in direct contact with the CoV-2 RBD residues. Lastly, we found several mutants that are located distal to the ACE2/S-protein binding site and do not mediate direct or indirect contacts with CoV-2 RBD. We classified these variants as “not relevant” in the context of our analysis. The enhancing variants were enriched in the mutagenesis screen (Procko, 2020), while those ACE2 variants predicted to disrupt or weaken the ACE2/S-protein interactions were depleted (Procko, 2020).

Polymorphic variants mapped onto the ACE2 structure remarkably segregate into two distinct clusters at the ACE2/CoV-2 RBD interface (**Figure 2**). Enhancing variants cluster to the ACE2 surface most proximal to the receptor-binding ridge of CoV-2 RBD (**Figure 3a**) whereas the majority of the disrupting variants reside centrally on the two major ACE2 α-helices that substantially contribute to the buried surface area at the interface (**Figure 4a**). Interestingly, the loop conformation in the receptor-binding ridge differs significantly in SARS-CoV-2 from that of SARS-CoV owing to the presence of bulky residues (V483 and E484) in the loop (Shang et al., 2020). This feature allows the CoV-2 loop to extend further towards ACE2 establishing more extensive contacts with the receptor. Hence, natural ACE2 variants in this region could be exploited by the CoV-2 loop, increasing susceptibility to viral infection. In contrast, most interactions that CoV-2 makes with the core of the ACE2 interface are centered on two α-helices and are mostly not unique to CoV-2. They seem to encompass critical binding hotspots, discussed below, and thus centrally located polymorphic variants are more likely to reduce viral recognition.

By far the most frequent variant identified in our data, K26R (∼0.4% allele frequency), is predicted to enhance ACE2 affinity for SARS-CoV-2. Structural analysis of this polymorphism shows that K26 establishes polar contacts with the first mannose moiety of the ACE2 N90-linked glycan and likely stabilizes the position of the glycan relative to ACE2 (**Figure 3b**). As discussed above, the N90-linked glycan emerges as an important determinant of CoV-2 infectivity and may diminish ACE2 affinity for the RBD possibly through steric hindrance imposed by branching of the sugar modifications (Demogines et al., 2012). We predict that K26R would abrogate stabilizing polar contacts with N90, impairing coordination of the glycan (**Figure 3b**) and lead to an increase in the affinity of the virus to the ACE2 receptor. At the same time, R26 is now primed to establish backbone and side chain interactions with ACE2 D30 which then is poised to build a salt-bridge with CoV-2 RBD K417 (**Figure 3b**). The net effect of R26 polymorphism would then be the stabilization of core α-helices that increases ACE2 binding affinity to CoV-2 RBD at the cost of glycan rigidity. As discussed above, another enhancing variant, T92I, is structurally predicted to lead to similar effects by directly eliminating the N90-linked glycan.

Whereas the K26R variant stabilizes core ACE2 α-helical interactions, other naturally occurring polymorphic variants appear to locally destabilize or alter the N-terminus of ACE2 helix 1 (α1) conformation. The T27A mutant (**Figure 3c**) removes side chain-backbone and backbone-backbone interactions between T27 and E30 likely increasing the local dynamics of helix α1. This would allow the N-terminus of α1 to bend slightly and accommodate the unique CoV-2 RBD receptor binding-ridge loop that more intimately contacts ACE2 compared to its SARS-CoV counterpart. Another predicted effect of the T27A variant is increased hydrophobicity at the interface, which could contribute to an increase in binding affinity. Similar destabilizing patterns can be inferred for S19P and E23K (**Supplementary Table 4**). Thus, the local α1 N-terminal helical flexibility along with N90 glycan destabilization may help accommodate the protrusive CoV-2 RBD receptor-binding ridge to form more extensive contacts with ACE2 and facilitate viral entry specifically for this virus.

ACE2 polymorphic variants predicted to confer protection against the virus (**Supplementary Tables 2 and 4**) present compelling mechanistic explanations for how they may offer protection against the virus. The vast majority of disruptive variants map to the core α-helical bundle of ACE2 and to residues known to form contacts with the RBD (**Figure 4a**). There are two key hotspots in the α-helical bundle of the ACE2 interface that are important for CoV-2 RBD binding: K31 and K353 (**Figure 4a, right panel**). To enable interaction with the virus, these charged residues need to be accommodated in a largely hydrophobic environment at the binding interface and hence their neutralization is critical to the binding of coronavirus RBDs to human ACE2 (Li, 2008; Shang et al., 2020; Wu et al., 2012). A recent elegant study (Shang et al., 2020) showed that SARS-CoV-2 S-protein is more effective in neutralization of the lysine hotspots than SARS-CoV due to the presence of Q493 and L455 that stabilize K31, and N501 that stabilizes K353, (**Figure 4A, right panel**). Interestingly, K31R is one of the human ACE2 polymorphisms that we identified (**Supplementary Table 2**). Introduction of an arginine not only maintains the positive charge at position 31 but is also predicted to break an interaction with Q493 in the RBD (**Supplementary Figure 5a**) and destabilize the charge-neutralizing interaction with the virus. Thus, individuals carrying K31R ACE2 variants are predicted to be less prone to SARS-CoV-2 infection. While we did not identify any polymorphic variants at residue K353, we detected an ACE2 mutation that changes the identity of D38 residue (**Supplementary Table 2**), which forms an electrostatic interaction with K353 (**Supplementary Table 4**). This mutation (D38V) would compromise the neutralizing effect of the K353-D38 interaction at the interface and is predicted to significantly reduce binding affinity between the virus and the host receptor.

Another recurrent polymorphism in ACE2 maps to residue E35 and changes it to a lysine (**Supplementary Table 1**). E35 establishes a critical polar contact with SARS-CoV-2 S-protein residue Q493, which is predicted to be attenuated in the presence of the positively charged lysine (**Figure 4B**). Interestingly, E35 is not conserved between SARS-CoV and SARS-CoV-2 S-proteins (Shang et al., 2020) and hence we predict that it could offer selective protection from the SARS-CoV-2 infection in individuals carrying this variant. Other variants found in our analysis, including H34R (**Figure 4C**) and D38V, similarly result in a loss of interface polar contacts which are predicted to reduce ACE2 affinity for the viral RBD domain. Another interesting polymorphism at position 83 results in Y83H alteration. Residue Y83 underlies a hydrophobic pocket into which F486 from SARS-CoV-2 RBD is inserted (**Supplementary Figure 5b**). This is another unique interaction involving ACE2 and the SARS-CoV-2 RBD F486 that is absent in SARS-CoV RBD where, the equivalent residue is a leucine (Shang et al., 2020). The polymorphism that replaces Y83 with a polar histidine will compromise the hydrophobic character of this unique pocket in addition to removing a polar contact with N487 (**Supplementary Figure 5b**), potentially offering selective protection from the SARS CoV-2 infections.

## Discussion

The host-virus evolutionary arms race over time leads to natural selection that alters both the host and the viral proteins allowing both to increase their fitness (Daugherty and Malik, 2012). In this context multiple studies have analyzed and identified the origin, evolution and successful adaption of the SARS coronaviruses as human pathogens (Andersen et al., 2020; Guo et al., 2020). Viral genome sequencing and analysis has identified bats as the most likely natural host of origin for both SARS-CoV and the recent SARS-CoV-2 (Guo et al., 2020). In particular, several studies have focused on the viral S-protein RBD that interacts with its host ACE2 receptor and identified key changes between the bat CoVs and other suspected intermediary host CoVs found in the civet and pangolin (Andersen et al., 2020; Chen et al., 2020; Shang et al., 2020; Walls et al., 2020; Wrapp et al., 2020; Yan et al., 2020). These studies have identified S-protein changes that have rendered the human cells permissive to the SARS-CoV and SARS-CoV-2 infection (Chen et al., 2020; Shang et al., 2020; Walls et al., 2020; Wrapp et al., 2020; Yan et al., 2020).

Thus far, the role of variations in human ACE2 receptor in susceptibility to both SARS CoVs had not been comprehensively examined. A recent study analyzed a limited ACE2 population variation data set and concluded that these polymorphisms did not confer resistance to the virus (Cao et al., 2020a). In this study, we have examined human ACE2 variation data compiled from multiple data sets and identified polymorphisms that will either likely render individuals more susceptible to the SARS-CoV-2 or protect them from the virus. Using structural predictions based on published protein structures and data from an elegant mutagenesis screen that used deep sequencing to assess enrichment or depletion of S-protein binding ACE2 variants, we classified the variants identified in this study for the effects on susceptibility to SARS-CoV (Procko, 2020; Shang et al., 2020; Walls et al., 2020; Wrapp et al., 2020; Yan et al., 2020). In particular, human ACE2 variants K26R, S16P, T27A, K31R, H34R, E35K, E37K, D38V, N51S, N64K, K68E, F72V, T921, Q102P, G326E, G352V, D355N, H378R, Q388L, and D509Y are predicted to increase the susceptibility of the individuals carrying these variations. It is interesting to note that the T921I ACE2 variant is part of the consensus NxS/T N-glycosylation motif and is predicted to abolish glycosylation of the conserved N90 residue. Our structural investigation suggests that this mutation will favor improved viral S-protein binding. A previous study showed that the ACE2 N90 renders human cells resistant to Civet CoV (Li et al., 2005b). Recently, N90 and T92 ACE2 mutations were enriched in a screen for CoV-2 S-protein binding (Procko, 2020). Taken together, these observations suggest that N90 and T92 are critical ACE2 residues that confer protection and are CoV host modifiers. We also found variants K31R, E35K, E37K, D38V, N33I, H34R, Q388L and Y83H in ACE2 that are predicted to show decreased binding to SARS-CoV-2 S-protein and thus protect individuals corresponding to these genotypes.

Overall, we find the ACE2 population variants, that either increase or decrease susceptibility, to be rare, which is consistent with the overall low population ACE2 receptor polymorphisms (mean Fst 0.0167). Also, we did not observe significant differences in ACE2 variant allele frequency among population groups. The variant alleles also did not show discernable gender distribution differences, even though ACE2 is a X-linked gene. The SARS-CoV infections and its deadly effects in humans are more recent and thus the pathogenic and protective variants have not been subject to purifying selection and therefore the variants we observe are predictably rare.

The expression levels of ACE2 and its variants in appropriate host tissue may modulate the deleterious effect of the virus. To further understand the importance of the ACE2 variants in susceptibility, it will be important to correlate clinical outcomes with ACE2 genotypes at population scale. The extremes in COVID-19 clinical symptoms reported ranging from asymptomatic infected adult individuals to those that show acute respiratory syndrome leading to death (Cao et al., 2020b; Cascella et al., 2020; Yuen et al., 2020), suggest a role for additional factors, including the role of innate and adaptive immunity, besides ACE2 variants in modifying disease outcomes.

Currently, there are no approved therapeutics for treating or preventing COVID-19 caused by the SARS-CoV-2. Therefore, development of therapeutics to treat patients and mitigate the COVID-19 pandemic is urgently needed (Cascella et al., 2020; Jiang, 2020). Several small molecules and neutralizing antibodies for treatment are in development (Li and De Clercq, 2020; Zhou et al., 2020b). Soluble ACE2 and ACE2-Fc fusion protein have been proposed as decoy SARS-CoV-2 receptor therapeutic (Hofmann et al., 2004; Kruse, 2020; Lei et al., 2020). Soluble ACE2, as a therapy for pulmonary arterial hypertension, has been shown to be safe in early in-human clinical studies (Guignabert et al., 2018; Haschke et al., 2013). A rationally designed, catalytically inactive, human ACE2 that carries one or more of the natural variants predicted to show improved binding to SARS viral S-protein RBD could be safely developed as a soluble protein with or without an Fc domain for treatment of COVID-19. Such a recombinant ACE2 protein can be engineered to create a pan-CoV neutralizing drug that is broad and can neutralize CoVs that may emerge during future epidemics.

## Methods

### Identification of ACE2 variation

We queried multiple genomic databases including gnomaAD (Karczewski et al., 2019) (https://gnomad.broadinstitute.org/), DicoverEHR (Dewey et al., 2016), RotterdamStudy (Ikram et al., 2017), ALSPAC (Fraser et al., 2013) and Asian specific databases which included GenomeAsia100k (GenomeAsia, 2019), HGDP (Bergstrom et al., 2020), TOMMO-3.5kjpnv2 (Tadaka et al., 2019) IndiGen (https://indigen.igib.in/) and Other aggregated data for ACE2 protein altering variations in populations groups across the world. The ACE2 genotypes in this study were from over 290,000 samples representing over 400 population groups across the world.

### Fst Analysis

To assess genetic variation in the coding region of ACE2, we calculated the fixation index (Fst) from 2,381 unrelated individuals across 26 populations in the 1000 Genomes Project Phase 3 and 57,783 female individuals across eight populations in gnomAD. For 1000 Genome data, we used the Weir and Cockerham (1984) method as implemented in vcftools (Version 0.1.17); the weighted Fst were calculated from 88 variants. For gnomAD (v2.1.1), because we only have access to the allele counts, we used the original formulation by Wright (1969) and reported the weighted mean Fst as described in Bhatia et al. (2013); 277 variants were used. Because Fst values vary based on variants used (Bhatia et al. 2013), we calculated the Fst in a set of randomly selected genes on the same chromosomes matched by the length decile to use for comparison. To assess if variants in the peptidase domain has lower genetic variation, we used the one-sided Wilcoxon rank-sum test to compare 15 variants in the peptidase domain against 50 variants outside. Variants with Fst < 1e-4 were removed as they were uninformative.

### ACE2 ortholog sequence analysis

A total of 295 Human ACE2 orthologs were obtained from NCBI (**Supplementary Table 3** for accession numbers). A snake ACE2 ortholog protein was obtained from the published Indian cobra genome (Suryamohan et al., 2020). Multiple sequence alignment of residues surrounding the ACE2 NxT/S motif was performed using MCoffee (www.tcoffee.org). Phylogenetic trees were constructed using the PhyML webserver (www.phylogeny.fr).

### Structural Analysis

Each identified variant was mapped, modeled, and analyzed in Pymol using the recently deposited crystal structures 6VW1 and 6LZG of human ACE2 bound to either chimeric SARS CoV-2 RBD (6VW1) or complete SARS CoV-2 RBD (6LZG).

## Acknowledgements

DD was supported by an F30 fellowship. Grant number: 1F30CA247147. AR and SCS were supported by SCELSE.

## Supplementary figures legend

**Figure S1.**
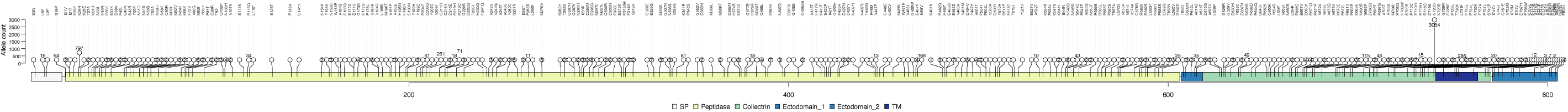
A cartoon of ACE2 protein showing protein altering polymorphic variants observed across the entire protein. Allele counts for each polymorphism is shown inside or above each circle. Empty circles indicate singletons.

**Figure S2.**
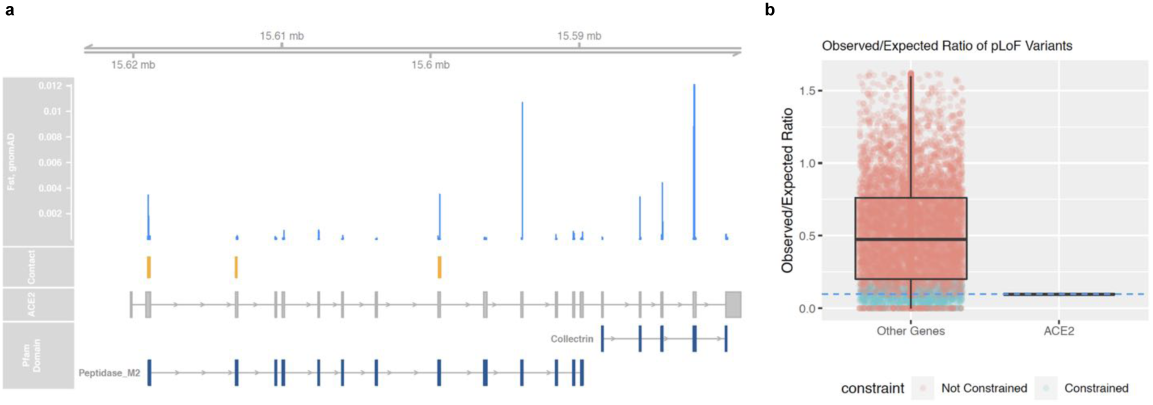
(a) Fst index of exonic variants of ACE2, calculated from 57,783 female individuals across eight populations in gnomAD. Canonical transcript of ACE2 (ENST00000427411) and two Pfam domains are shown along with the positions of known SARS-CoV-2 contact residues. Peptidase domain harbor variants with lower variation (Wilcox p=0.0656). (**b)** ACE2 is highly constrained (pLI=0.9977), with the observed-to-expected ratio of the number of pLoF variants of 0.0968, consistent with the constrained genes (highlighted in cyan).

**Figure S3.**
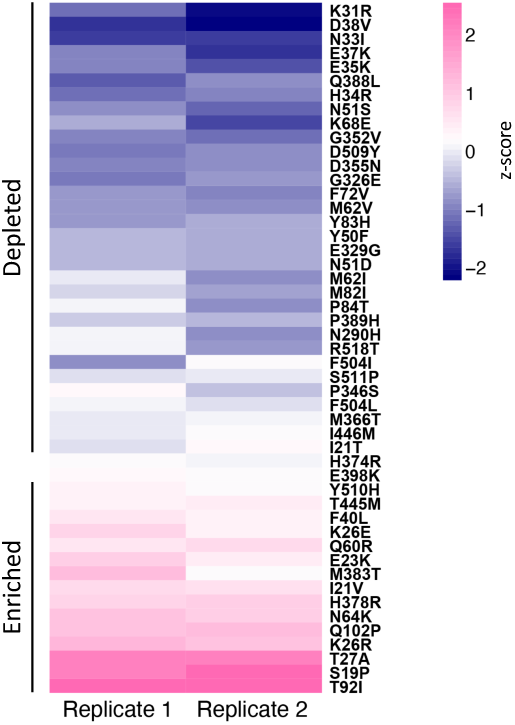
Heatmap showing human ACE2 polymorphism that map to the ACE2-RBD intraction region and the corresponding enrichment/depletion scores from a recent study by Procko 2020 (doi.org/10.1101/2020.03.16.994236)

**Figure S4.**
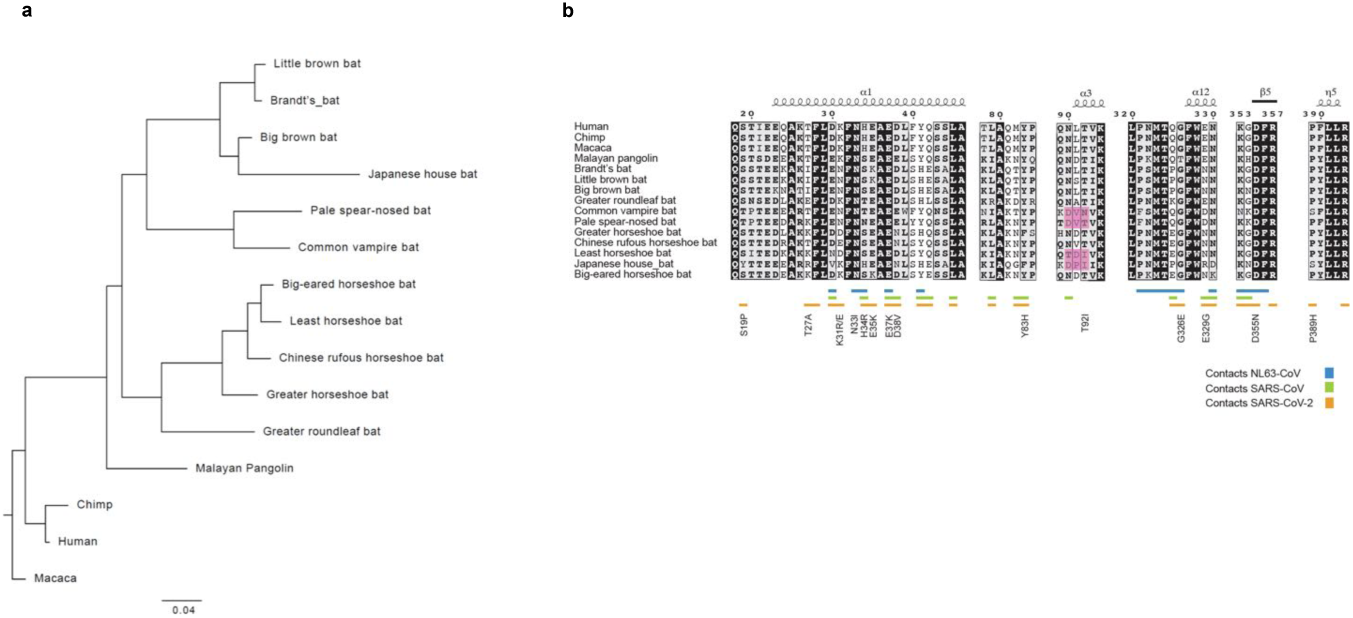
ACE2 sequence comparison. (**a**). Phylogenetic tree of ACE2 sequences from selected species, **(b)** Multiple sequence alignment of representative primate ACE2 sequences and ACE2 sequences of putative natural and intermediate reservoirs of coronaviruses. Pink boxes highlight species where the canonical NxT/S motif is absent or altered.

**Figure S5.**
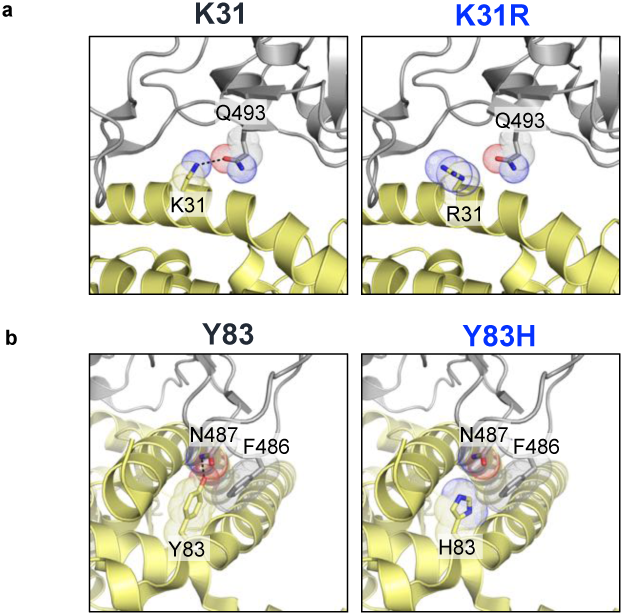
Structural basis for the destabilizing effects of K31R and Y83H human polymorphisms on the interactions with SARS-CoV-2. **(a)** Analysis of the polymorphism at K31 of ACE2 shows that its mutation to an arginine breaks an energetically favorable electrostatic interaction with Q493 of SARS-CoV-2 RBD (PDB: 6LZG). **(b)** Analysis of the effect of ACE2 Y83 mutation to histidine shows loss of polar contact with N487 as well as reduced hydrophobic packing with the unique SARS-CoV-2 RBD F486 (PDB: 6LZG).

