## Supplementary Tables 1-4 for "Human ACE2 receptor polymorphisms predict SARS-CoV-2 susceptibility"

**Supplementary Table 1 - Human ACE2 protein altering population variants**
**Supplementary Table 2 - ACE2 polymorphisms predicted to alter SARS-CoV/Cov-2 S-protein binding**
**Supplementary Table 3 - Conservation of N90 glycosylation motif in annotated jawed vertebrate ACE2 orthologs**
**Supplementary Table 4 - Structure evaluation findings**

| | A | B | C | D | E | F | G | H | I |
|---|---|---|---|---|---|---|---|---|---|
| 1 | **Supplementary Table 1 - Human ACE2 protein altering population variants** | | | | | | | | |
| 2 | **Position** | **Mutation** | **Protein Consequence** | **Allele Count** | **gnomAD Allele Frequency** | **Primary Source** | **Additional Sources** | **Mean Procko Score** | **Contact Residue** |
| 3 | 3 | S3N | p.Ser3Asn | 1 | 5.83625E-06 | gnomAD | AsianSpecificDB | | |
| 4 | 8 | L8F | p.Leu8Phe | 16 | 8.05814E-05 | gnomAD | Other | | |
| 5 | 9 | L9P | p.Leu9Pro | 1 | 5.62943E-06 | gnomAD | Other | | |
| 6 | 19 | S19P | p.Ser19Pro | 64 | 0.000312919 | gnomAD | Other | 2.327818365 | |
| 7 | 21 | I21V | p.Ile21Val | 2 | 1.0927E-05 | gnomAD | Other | 0.697012397 | |
| 8 | 21 | I21T | p.Ile21Thr | 1 | 5.46254E-06 | gnomAD | | 0.072108031 | |
| 9 | 23 | E23K | p.Glu23Lys | 1 | 5.45777E-06 | gnomAD | | 0.647777027 | |
| 10 | 26 | K26R | p.Lys26Arg | 797 | 0.003883296 | gnomAD | AsianSpecificDB,UKBio | 1.229246696 | |
| 11 | 26 | K26E | p.Lys26Glu | 1 | 5.45476E-06 | gnomAD | | 0.573492928 | |
| 12 | 27 | T27A | p.Thr27Ala | 2 | 1.09099E-05 | gnomAD | | 2.113424949 | Y |
| 13 | 31 | K31R | p.Lys31Arg | 1 | 0 | AsianSpecif | AsianSpecificDB | -1.609477184 | Y |
| 14 | 33 | N33I | p.Asn33Ile | 1 | 0 | AsianSpecif | AsianSpecificDB | -1.568303739 | |
| 15 | 34 | H34R | p.His34Arg | 1 | 0 | AsianSpecif | AsianSpecificDB | -1.076031319 | Y |
| 16 | 35 | E35K | p.Glu35Lys | 3 | 1.636E-05 | gnomAD | | -1.199253155 | Y |
| 17 | 37 | E37K | p.Glu37Lys | 8 | 3.89708E-05 | gnomAD | AsianSpecificDB | -1.349986562 | Y |
| 18 | 38 | D38V | p.Asp38Val | 1 | 0 | Other | | -1.963734719 | Y |
| 19 | 40 | F40L | p.Phe40Leu | 2 | 3.0862E-05 | Gnomad | | 0.449051719 | |
| 20 | 43 | S43N | p.Ser43Asn | 2 | 0 | Other | | | |
| 21 | 43 | S43R | p.Ser43Arg | 1 | 5.46185E-06 | gnomAD | | | |
| 22 | 50 | Y50F | p.Tyr50Phe | 1 | 5.4817E-06 | gnomAD | | -0.535360358 | |
| 23 | 51 | N51D | p.Asn51Asp | 1 | 5.48501E-06 | gnomAD | | -0.500865398 | |
| 24 | 51 | N51S | p.Asn51Ser | 1 | 5.48781E-06 | gnomAD | | -1.052778389 | |
| 25 | 53 | N53S | p.Asn53Ser | 1 | 0 | AsianSpecif | AsianSpecificDB | | |
| 26 | 55 | T55A | p.Thr55Ala | 1 | 5.52312E-06 | gnomAD | | | |
| 27 | 58 | N58H | p.Asn58His | 2 | 1.11702E-05 | gnomAD | | | |
| 28 | 58 | N58K | p.Asn58Lys | 2 | 1.11867E-05 | gnomAD | | | |
| 29 | 60 | Q60R | p.Gln60Arg | 2 | 1.12455E-05 | gnomAD | | 0.602848439 | |
| 30 | 62 | M62I | p.Met62Ile | 1 | 0 | AsianSpecif | AsianSpecificDB | -0.433221503 | |

| | A | B | C | D | E | F | G | H | I |
|---|---|---|---|---|---|---|---|---|---|
| 2 | **Position** | **Mutation** | **Protein Consequence** | **Allele Count** | **gnomAD Allele Frequency** | **Primary Source** | **Additional Sources** | **Mean Procko Score** | **Contact Residue** |
| 31 | 62 | M62V | p.Met62Val | 1 | 5.66646E-06 | gnomAD | AsianSpecificDB | -0.782795497 | |
| 32 | 64 | N64K | p.Asn64Lys | 3 | 1.4664E-05 | gnomAD | | 1.047209649 | |
| 33 | 68 | K68E | p.Lys68Glu | 2 | 1.09457E-05 | gnomAD | AsianSpecificDB | -1.052667031 | |
| 34 | 72 | F72V | p.Phe72Val | 1 | 5.47411E-06 | gnomAD | | -0.88673533 | |
| 35 | 80 | A80G | p.Ala80Gly | 1 | 0 | AsianSpecif | AsianSpecificDB | | |
| 36 | 82 | M82I | p.Met82Ile | 5 | 2.44178E-05 | gnomAD | | -0.428501783 | Y |
| 37 | 83 | Y83H | p.Tyr83His | 2 | 0 | SAS(1) | AsianSpecificDB | -0.667134906 | Y |
| 38 | 84 | P84T | p.Pro84Thr | 1 | 5.4707E-06 | gnomAD | | -0.419131731 | |
| 39 | 86 | Q86R | p.Gln86Arg | 2 | 1.09411E-05 | gnomAD | Other | | |
| 40 | 92 | T92I | p.Thr92Ile | 2 | 1.09557E-05 | gnomAD | | 2.534286782 | |
| 41 | 102 | Q102P | p.Gln102Pro | 3 | 1.47451E-05 | gnomAD | | 1.191328285 | |
| 42 | 103 | N103H | p.Asn103His | 4 | 1.96661E-05 | gnomAD | | | |
| 43 | 107 | V107A | p.Val107Ala | 2 | 1.10552E-05 | gnomAD | Other | | |
| 44 | 113 | S113N | p.Ser113Asn | 1 | 0 | Other | | | |
| 45 | 115 | R115Q | p.Arg115Gln | 34 | 0.000170308 | gnomAD | Other | | |
| 46 | 128 | S128T | p.Ser128Thr | 1 | 5.73809E-06 | gnomAD | | | |
| 47 | 138 | P138A | p.Pro138Ala | 1 | 5.73283E-06 | gnomAD | | | |
| 48 | 141 | C141Y | p.Cys141Tyr | 1 | 5.8307E-06 | gnomAD | | | |
| 49 | 154 | N154K | p.Asn154Lys | 2 | 1.09536E-05 | gnomAD | | | |
| 50 | 158 | Y158H | p.Tyr158His | 1 | 0 | Other | | | |
| 51 | 159 | N159S | p.Asn159Ser | 3 | 1.63682E-05 | gnomAD | AsianSpecificDB,UKBio | | |
| 52 | 163 | W163R | p.Trp163Arg | 1 | 5.45557E-06 | gnomAD | | | |
| 53 | 164 | A164S | p.Ala164Ser | 1 | 5.45461E-06 | gnomAD | | | |
| 54 | 166 | E166Q | p.Glu166Gln | 1 | 5.4536E-06 | gnomAD | | | |
| 55 | 171 | E171D | p.Glu171Asp | 1 | 4.55519E-05 | gnomAD | | | |
| 56 | 171 | E171V | p.Glu171Val | 1 | 5.45411E-06 | gnomAD | | | |
| 57 | 173 | G173S | p.Gly173Ser | 4 | 2.18172E-05 | gnomAD | | | |
| 58 | 177 | R177S | p.Arg177Ser | 1 | 0 | AsianSpecif | AsianSpecificDB | | |
| 59 | 178 | P178L | p.Pro178Leu | 1 | 5.455E-06 | gnomAD | | | |
| 60 | 184 | V184A | p.Val184Ala | 8 | 4.36503E-05 | gnomAD | AsianSpecificDB,UKBio | | |
| 61 | 184 | V184G | p.Val184Gly | 1 | 4.58358E-05 | gnomAD | | | |

| | A | B | C | D | E | F | G | H | I |
|---|---|---|---|---|---|---|---|---|---|
| 2 | **Position** | **Mutation** | **Protein Consequence** | **Allele Count** | **gnomAD Allele Frequency** | **Primary Source** | **Additional Sources** | **Mean Procko Score** | **Contact Residue** |
| 62 | 186 | L186S | p.Leu186Ser | 1 | 5.4615E-06 | gnomAD | | | |
| 63 | 190 | M190T | p.Met190Thr | 1 | 5.46591E-06 | gnomAD | | | |
| 64 | 191 | A191P | p.Ala191Pro | 1 | 5.46819E-06 | gnomAD | | | |
| 65 | 193 | A193E | p.Ala193Glu | 3 | 1.64126E-05 | gnomAD | | | |
| 66 | 195 | H195N | p.His195Asn | 1 | 5.47627E-06 | gnomAD | AsianSpecificDB | | |
| 67 | 195 | H195Y | p.His195Tyr | 1 | 5.47627E-06 | gnomAD | | | |
| 68 | 198 | D198N | p.Asp198Asn | 1 | 5.88166E-06 | gnomAD | | | |
| 69 | 199 | Y199H | p.Tyr199His | 1 | 0 | AsianSpecif | AsianSpecificDB | | |
| 70 | 199 | Y199C | p.Tyr199Cys | 3 | 1.68777E-05 | gnomAD | AsianSpecificDB | | |
| 71 | 204 | R204T | p.Arg204Thr | 1 | 5.48152E-06 | gnomAD | AsianSpecificDB | | |
| 72 | 206 | D206G | p.Asp206Gly | 61 | 0.000299999 | gnomAD | Other | | |
| 73 | 207 | Y207C | p.Tyr207Cys | 1 | 5.46866E-06 | gnomAD | | | |
| 74 | 209 | V209I | p.Val209Ile | 1 | 5.46735E-06 | gnomAD | | | |
| 75 | 211 | G211R | p.Gly211Arg | 261 | 0.001279889 | gnomAD | AsianSpecificDB,UKBio | | |
| 76 | 216 | D216Y | p.Asp216Tyr | 1 | 5.46015E-06 | gnomAD | Other | | |
| 77 | 216 | D216E | p.Asp216Glu | 3 | 1.47049E-05 | gnomAD | | | |
| 78 | 219 | R219H | p.Arg219His | 18 | 9.83128E-05 | gnomAD | AsianSpecificDB,UKBio | | |
| 79 | 219 | R219C | p.Arg219Cys | 71 | 0.000348058 | gnomAD | Other | | |
| 80 | 220 | G220S | p.Gly220Ser | 3 | 1.63894E-05 | gnomAD | AsianSpecificDB | | |
| 81 | 225 | D225G | p.Asp225Gly | 1 | 0 | AsianSpecif | AsianSpecificDB | | |
| 82 | 229 | T229I | p.Thr229Ile | 1 | 5.4787E-06 | gnomAD | | | |
| 83 | 239 | H239Q | p.His239Gln | 1 | 5.85967E-06 | gnomAD | | | |
| 84 | 241 | H241Q | p.His241Gln | 1 | 0 | Other | | | |
| 85 | 242 | A242V | p.Ala242Val | 3 | 1.7254E-05 | gnomAD | | | |
| 86 | 246 | A246T | p.Ala246Thr | 2 | 0 | AsianSpecif | AsianSpecificDB | | |
| 87 | 246 | A246S | p.Ala246Ser | 1 | 5.68311E-06 | gnomAD | | | |
| 88 | 252 | Y252C | p.Tyr252Cys | 2 | 1.12827E-05 | gnomAD | AsianSpecificDB | | |
| 89 | 257 | S257N | p.Ser257Asn | 5 | 2.57054E-05 | gnomAD | AsianSpecificDB | | |
| 90 | 259 | I259T | p.Ile259Thr | 2 | 1.03069E-05 | gnomAD | | | |
| 91 | 263 | P263S | p.Pro263Ser | 11 | 5.78415E-05 | gnomAD | Other | | |
| 92 | 266 | L266F | p.Leu266Phe | 1 | 0 | Other | | | |

| | A | B | C | D | E | F | G | H | I |
|---|---|---|---|---|---|---|---|---|---|
| 2 | Position | Mutation | Protein Consequence | Allele Count | gnomAD Allele Frequency | Primary Source | Additional Sources | Mean Procko Score | Contact Residue |
| 93 | 270 | M270V | p.Met270Val | 5 | 2.98299E-05 | gnomAD | AsianSpecificDB,UKBio | | |
| 94 | 280 | S280Y | p.Ser280Tyr | 1 | 5.65141E-06 | gnomAD | | | |
| 95 | 282 | T282S | p.Thr282Ser | 2 | 0 | AsianSpecif | AsianSpecificDB | | |
| 96 | 287 | Q287R | p.Gln287Arg | 1 | 5.66046E-06 | gnomAD | | | |
| 97 | 287 | Q287K | p.Gln287Lys | 1 | 5.65384E-06 | gnomAD | | | |
| 98 | 290 | N290H | p.Asn290His | 2 | 1.13323E-05 | gnomAD | | -0.382086808 | |
| 99 | 291 | I291K | p.Ile291Lys | 3 | 1.70594E-05 | gnomAD | Other | | |
| 100 | 292 | D292N | p.Asp292Asn | 2 | 1.13967E-05 | gnomAD | | | |
| 101 | 295 | D295G | p.Asp295Gly | 8 | 4.6594E-05 | gnomAD | Other | | |
| 102 | 297 | M297I | p.Met297Ile | 1 | 5.86304E-06 | gnomAD | | | |
| 103 | 297 | M297L | p.Met297Leu | 1 | 5.85144E-06 | gnomAD | | | |
| 104 | 300 | Q300R | p.Gln300Arg | 2 | 1.18694E-05 | gnomAD | | | |
| 105 | 303 | D303N | p.Asp303Asn | 2 | 1.20266E-05 | gnomAD | AsianSpecificDB | | |
| 106 | 308 | F308L | p.Phe308Leu | 1 | 5.73365E-06 | gnomAD | | | |
| 107 | 312 | E312K | p.Glu312Lys | 2 | 1.13409E-05 | gnomAD | | | |
| 108 | 314 | F314S | p.Phe314Ser | 2 | 0 | Other | | | |
| 109 | 318 | V318A | p.Val318Ala | 2 | 0 | AsianSpecif | AsianSpecificDB | | |
| 110 | 326 | G326E | p.Gly326Glu | 1 | 5.51675E-06 | gnomAD | | -0.90546886 | Y |
| 111 | 329 | E329G | p.Glu329Gly | 7 | 3.44298E-05 | gnomAD | Other | -0.506466064 | Y |
| 112 | 332 | M332L | p.Met332Leu | 3 | 1.65432E-05 | gnomAD | | | |
| 113 | 337 | G337R | p.Gly337Arg | 1 | 5.53168E-06 | gnomAD | | | |
| 114 | 338 | N338S | p.Asn338Ser | 3 | 1.65937E-05 | gnomAD | | | |
| 115 | 339 | V339G | p.Val339Gly | 1 | 5.53661E-06 | gnomAD | | | |
| 116 | 341 | K341R | p.Lys341Arg | 81 | 0.000400192 | gnomAD | AsianSpecificDB,UKBio | | |
| 117 | 346 | P346S | p.Pro346Ser | 1 | 5.59945E-06 | gnomAD | | -0.066906284 | |
| 118 | 352 | G352V | p.Gly352Val | 1 | 5.75218E-06 | gnomAD | | -1.025319393 | |
| 119 | 355 | D355N | p.Asp355Asn | 2 | 1.17433E-05 | gnomAD | Other | -0.911052386 | |
| 120 | 360 | M360L | p.Met360Leu | 1 | 5.53836E-06 | gnomAD | | | |
| 121 | 366 | M366T | p.Met366Thr | 2 | 1.09845E-05 | gnomAD | | 0.034196848 | |
| 122 | 368 | D368N | p.Asp368Asn | 1 | 5.49125E-06 | gnomAD | AsianSpecificDB | | |
| 123 | 374 | H374R | p.His374Arg | 1 | 5.47528E-06 | gnomAD | | 0.110711661 | |

| | A | B | C | D | E | F | G | H | I |
|---|---|---|---|---|---|---|---|---|---|
| 2 | **Position** | **Mutation** | **Protein Consequence** | **Allele Count** | **gnomAD Allele Frequency** | **Primary Source** | **Additional Sources** | **Mean Procko Score** | **Contact Residue** |
| 124 | 375 | E375D | p.Glu375Asp | 3 | 1.64213E-05 | gnomAD | | | |
| 125 | 377 | G377E | p.Gly377Glu | 1 | 5.47345E-06 | gnomAD | | | |
| 126 | 378 | H378R | p.His378Arg | 18 | 8.79104E-05 | gnomAD | Other | 0.83154494 | |
| 127 | 383 | M383T | p.Met383Thr | 1 | 0 | AsianSpecif | AsianSpecificDB | 0.676250072 | |
| 128 | 388 | Q388L | p.Gln388Leu | 4 | 2.18607E-05 | gnomAD | | -1.084467232 | |
| 129 | 389 | P389H | p.Pro389His | 7 | 3.82576E-05 | gnomAD | | -0.400730258 | |
| 130 | 397 | N397D | p.Asn397Asp | 3 | 1.46412E-05 | gnomAD | | | |
| 131 | 398 | E398K | p.Glu398Lys | 1 | 5.46224E-06 | gnomAD | | 0.205797456 | |
| 132 | 405 | G405E | p.Gly405Glu | 1 | 5.46072E-06 | gnomAD | | | |
| 133 | 413 | A413T | p.Ala413Thr | 1 | 0 | Other | | | |
| 134 | 417 | H417R | p.His417Arg | 1 | 0 | AsianSpecif | AsianSpecificDB | | |
| 135 | 419 | K419T | p.Lys419Thr | 1 | 5.46263E-06 | gnomAD | | | |
| 136 | 420 | S420P | p.Ser420Pro | 1 | 4.54876E-05 | gnomAD | | | |
| 137 | 421 | I421T | p.Ile421Thr | 1 | 5.46379E-06 | gnomAD | | | |
| 138 | 426 | P426A | p.Pro426Ala | 1 | 5.46866E-06 | gnomAD | | | |
| 139 | 427 | D427N | p.Asp427Asn | 2 | 0 | Other | | | |
| 140 | 427 | D427Y | p.Asp427Tyr | 2 | 1.0948E-05 | gnomAD | | | |
| 141 | 437 | N437H | p.Asn437His | 1 | 0 | AsianSpecif | AsianSpecificDB | | |
| 142 | 437 | N437S | p.Asn437Ser | 1 | 0 | AsianSpecif | AsianSpecificDB | | |
| 143 | 445 | T445M | p.Thr445Met | 1 | 5.86937E-06 | gnomAD | | 0.422616407 | |
| 144 | 446 | I446M | p.Ile446Met | 1 | 4.53968E-05 | gnomAD | | 0.059171832 | |
| 145 | 447 | V447F | p.Val447Phe | 13 | 6.68439E-05 | gnomAD | | | |
| 146 | 448 | G448E | p.Gly448Glu | 1 | 5.77481E-06 | gnomAD | | | |
| 147 | 450 | L450V | p.Leu450Val | 1 | 5.71984E-06 | gnomAD | | | |
| 148 | 455 | M455I | p.Met455Ile | 1 | 5.65163E-06 | gnomAD | | | |
| 149 | 461 | W461R | p.Trp461Arg | 1 | 5.59851E-06 | gnomAD | | | |
| 150 | 463 | V463I | p.Val463Ile | 1 | 5.57479E-06 | gnomAD | | | |
| 151 | 466 | G466W | p.Gly466Trp | 1 | 5.57336E-06 | gnomAD | | | |
| 152 | 467 | E467K | p.Glu467Lys | 4 | 2.24252E-05 | gnomAD | | | |
| 153 | 468 | I468V | p.Ile468Val | 168 | 0.000838441 | gnomAD | AsianSpecificDB,UKBio | | |
| 154 | 481 | K481N | p.Lys481Asn | 1 | 0 | Other | | | |

| | A | B | C | D | E | F | G | H | I |
|---|---|---|---|---|---|---|---|---|---|
| 2 | **Position** | **Mutation** | **Protein Consequence** | **Allele Count** | **gnomAD Allele Frequency** | **Primary Source** | **Additional Sources** | **Mean Procko Score** | **Contact Residue** |
| 155 | 482 | R482Q | p.Arg482Gln | 3 | 1.7609E-05 | gnomAD | Other | | |
| 156 | 483 | E483Q | p.Glu483Gln | 1 | 0 | AsianSpecif | AsianSpecificDB | | |
| 157 | 483 | E483D | p.Glu483Asp | 7 | 4.63908E-05 | gnomAD | | | |
| 158 | 488 | V488A | p.Val488Ala | 1 | 6.32495E-06 | gnomAD | | | |
| 159 | 491 | V491M | p.Val491Met | 1 | 6.28828E-06 | gnomAD | | | |
| 160 | 494 | D494V | p.Asp494Val | 8 | 4.9578E-05 | gnomAD | Other | | |
| 161 | 497 | Y497H | p.Tyr497His | 1 | 0 | AsianSpecif | AsianSpecificDB | | |
| 162 | 501 | A501T | p.Ala501Thr | 4 | 2.21659E-05 | gnomAD | AsianSpecificDB,UKBio | | |
| 163 | 504 | F504I | p.Phe504Ile | 2 | 1.26824E-05 | gnomAD | | -0.33180346 | |
| 164 | 504 | F504L | p.Phe504Leu | 1 | 4.6307E-05 | gnomAD | | -0.035560813 | |
| 165 | 506 | V506A | p.Val506Ala | 1 | 6.56052E-06 | gnomAD | | | |
| 166 | 509 | D509Y | p.Asp509Tyr | 1 | 0 | Other | | -0.942921896 | |
| 167 | 510 | Y510H | p.Tyr510His | 1 | 6.86304E-06 | gnomAD | | 0.246017376 | |
| 168 | 511 | S511P | p.Ser511Pro | 1 | 0 | AsianSpecif | AsianSpecificDB | -0.096523335 | |
| 169 | 518 | R518T | p.Arg518Thr | 1 | 5.4904E-06 | gnomAD | | -0.34082089 | |
| 170 | 519 | T519I | p.Thr519Ile | 1 | 5.48276E-06 | gnomAD | | | |
| 171 | 521 | Y521H | p.Tyr521His | 1 | 5.47312E-06 | gnomAD | | | |
| 172 | 527 | E527V | p.Glu527Val | 1 | 0 | AsianSpecif | AsianSpecificDB | | |
| 173 | 532 | A532T | p.Ala532Thr | 10 | 5.46305E-05 | gnomAD | AsianSpecificDB | | |
| 174 | 534 | K534R | p.Lys534Arg | 1 | 5.46138E-06 | gnomAD | | | |
| 175 | 538 | P538L | p.Pro538Leu | 1 | 5.46379E-06 | gnomAD | | | |
| 176 | 541 | K541N | p.Lys541Asn | 1 | 0 | Other | | | |
| 177 | 541 | K541I | p.Lys541Ile | 2 | 9.75895E-06 | gnomAD | | | |
| 178 | 544 | I544N | p.Ile544Asn | 1 | 5.47714E-06 | gnomAD | | | |
| 179 | 546 | N546D | p.Asn546Asp | 8 | 3.91654E-05 | gnomAD | | | |
| 180 | 546 | N546S | p.Asn546Ser | 1 | 5.48585E-06 | gnomAD | | | |
| 181 | 547 | S547C | p.Ser547Cys | 43 | 0.00021059 | gnomAD | | | |
| 182 | 547 | S547F | p.Ser547Phe | 1 | 5.49058E-06 | gnomAD | | | |
| 183 | 550 | A550G | p.Ala550Gly | 2 | 0 | Other | | | |
| 184 | 553 | K553T | p.Lys553Thr | 2 | 9.84518E-06 | gnomAD | | | |
| 185 | 559 | R559S | p.Arg559Ser | 1 | 5.6744E-06 | gnomAD | | | |

| | A | B | C | D | E | F | G | H | I |
|---|---|---|---|---|---|---|---|---|---|
| 2 | **Position** | **Mutation** | **Protein Consequence** | **Allele Count** | **gnomAD Allele Frequency** | **Primary Source** | **Additional Sources** | **Mean Procko Score** | **Contact Residue** |
| 186 | 563 | S563L | p.Ser563Leu | 1 | 5.6211E-06 | gnomAD | | | |
| 187 | 565 | P565T | p.Pro565Thr | 1 | 0 | AsianSpecif | AsianSpecificDB | | |
| 188 | 565 | P565S | p.Pro565Ser | 1 | 5.57588E-06 | gnomAD | | | |
| 189 | 567 | T567A | p.Thr567Ala | 1 | 5.55222E-06 | gnomAD | | | |
| 190 | 570 | L570S | p.Leu570Ser | 1 | 4.59876E-05 | gnomAD | Other | | |
| 191 | 573 | V573A | p.Val573Ala | 2 | 1.09806E-05 | gnomAD | Other | | |
| 192 | 574 | V574I | p.Val574Ile | 1 | 5.49082E-06 | gnomAD | AsianSpecificDB | | |
| 193 | 574 | V574A | p.Val574Ala | 1 | 0 | AsianSpecif | AsianSpecificDB | | |
| 194 | 575 | G575R | p.Gly575Arg | 1 | 5.48799E-06 | gnomAD | | | |
| 195 | 582 | R582K | p.Arg582Lys | 3 | 1.64348E-05 | gnomAD | Other | | |
| 196 | 582 | R582S | p.Arg582Ser | 3 | 1.46802E-05 | gnomAD | | | |
| 197 | 585 | L585P | p.Leu585Pro | 1 | 5.47948E-06 | gnomAD | | | |
| 198 | 586 | N586Y | p.Asn586Tyr | 2 | 9.78852E-06 | gnomAD | | | |
| 199 | 588 | F588S | p.Phe588Ser | 1 | 5.48252E-06 | gnomAD | | | |
| 200 | 589 | E589G | p.Glu589Gly | 1 | 5.48477E-06 | gnomAD | | | |
| 201 | 593 | T593N | p.Thr593Asn | 3 | 1.47442E-05 | gnomAD | | | |
| 202 | 595 | L595V | p.Leu595Val | 3 | 1.65411E-05 | gnomAD | | | |
| 203 | 597 | D597E | p.Asp597Glu | 25 | 0.000123366 | gnomAD | AsianSpecificDB,UKBio | | |
| 204 | 608 | T608I | p.Thr608Ile | 1 | 5.67617E-06 | gnomAD | | | |
| 205 | 609 | D609N | p.Asp609Asn | 3 | 1.71863E-05 | gnomAD | Other | | |
| 206 | 612 | P612L | p.Pro612Leu | 2 | 0 | AsianSpecif | AsianSpecificDB | | |
| 207 | 614 | A614S | p.Ala614Ser | 35 | 0.00017115 | gnomAD | Other | | |
| 208 | 614 | A614T | p.Ala614Thr | 1 | 0 | Other | | | |
| 209 | 615 | D615G | p.Asp615Gly | 5 | 2.73928E-05 | gnomAD | | | |
| 210 | 627 | A627V | p.Ala627Val | 2 | 1.0948E-05 | gnomAD | | | |
| 211 | 628 | L628P | p.Leu628Pro | 1 | 0 | Other | | | |
| 212 | 629 | G629R | p.Gly629Arg | 1 | 0 | AsianSpecif | AsianSpecificDB | | |
| 213 | 629 | G629V | p.Gly629Val | 1 | 5.47411E-06 | gnomAD | | | |
| 214 | 630 | D630H | p.Asp630His | 3 | 1.64245E-05 | gnomAD | | | |
| 215 | 633 | Y633C | p.Tyr633Cys | 1 | 5.93384E-06 | gnomAD | | | |
| 216 | 637 | D637N | p.Asp637Asn | 3 | 0 | Other | | | |

| | A | B | C | D | E | F | G | H | I |
|---|---|---|---|---|---|---|---|---|---|
| 2 | **Position** | **Mutation** | **Protein Consequence** | **Allele Count** | **gnomAD Allele Frequency** | **Primary Source** | **Additional Sources** | **Mean Procko Score** | **Contact Residue** |
| 217 | 638 | N638S | p.Asn638Ser | 49 | 0.000253484 | gnomAD | AsianSpecificDB,UKBio | | |
| 218 | 638 | N638D | p.Asn638Asp | 1 | 0 | AsianSpecif | AsianSpecificDB | | |
| 219 | 644 | R644Q | p.Arg644Gln | 1 | 0 | AsianSpecif | AsianSpecificDB | | |
| 220 | 646 | S646F | p.Ser646Phe | 1 | 0 | AsianSpecif | AsianSpecificDB | | |
| 221 | 652 | R652K | p.Arg652Lys | 1 | 5.75844E-06 | gnomAD | | | |
| 222 | 653 | Q653K | p.Gln653Lys | 1 | 0 | Other | | | |
| 223 | 654 | Y654S | p.Tyr654Ser | 1 | 4.55208E-05 | gnomAD | | | |
| 224 | 658 | V658I | p.Val658Ile | 1 | 5.95575E-06 | gnomAD | | | |
| 225 | 660 | N660S | p.Asn660Ser | 1 | 5.99797E-06 | gnomAD | | | |
| 226 | 664 | L664I | p.Leu664Ile | 2 | 0 | Other | | | |
| 227 | 665 | F665C | p.Phe665Cys | 1 | 0 | Other | | | |
| 228 | 667 | E667V | p.Glu667Val | 1 | 5.52077E-06 | gnomAD | Other | | |
| 229 | 668 | E668K | p.Glu668Lys | 4 | 2.19972E-05 | gnomAD | AsianSpecificDB | | |
| 230 | 671 | R671Q | p.Arg671Gln | 4 | 1.96187E-05 | gnomAD | | | |
| 231 | 671 | R671P | p.Arg671Pro | 1 | 4.61553E-05 | gnomAD | | | |
| 232 | 672 | V672A | p.Val672Ala | 1 | 5.48195E-06 | gnomAD | | | |
| 233 | 672 | V672L | p.Val672Leu | 2 | 1.09705E-05 | gnomAD | | | |
| 234 | 673 | A673G | p.Ala673Gly | 1 | 5.47921E-06 | gnomAD | | | |
| 235 | 673 | A673V | p.Ala673Val | 1 | 5.47921E-06 | gnomAD | | | |
| 236 | 676 | K676E | p.Lys676Glu | 1 | 0 | AsianSpecif | AsianSpecificDB | | |
| 237 | 677 | P677L | p.Pro677Leu | 1 | 5.47525E-06 | gnomAD | | | |
| 238 | 681 | F681V | p.Phe681Val | 1 | 5.47202E-06 | gnomAD | | | |
| 239 | 688 | P688R | p.Pro688Arg | 1 | 5.47282E-06 | gnomAD | | | |
| 240 | 689 | K689E | p.Lys689Glu | 3 | 1.64049E-05 | gnomAD | | | |
| 241 | 690 | N690S | p.Asn690Ser | 1 | 5.47211E-06 | gnomAD | | | |
| 242 | 692 | S692P | p.Ser692Pro | 115 | 0.000561883 | gnomAD | AsianSpecificDB,UKBio | | |
| 243 | 693 | D693G | p.Asp693Gly | 1 | 5.47217E-06 | gnomAD | | | |
| 244 | 696 | P696T | p.Pro696Thr | 2 | 1.09609E-05 | gnomAD | Other | | |
| 245 | 697 | R697G | p.Arg697Gly | 46 | 0.000252134 | gnomAD | AsianSpecificDB,UKBio | | |
| 246 | 703 | A703T | p.Ala703Thr | 1 | 0 | AsianSpecif | AsianSpecificDB | | |
| 247 | 703 | A703S | p.Ala703Ser | 3 | 1.65621E-05 | gnomAD | | | |

| | A | B | C | D | E | F | G | H | I |
|---|---|---|---|---|---|---|---|---|---|
| 2 | **Position** | **Mutation** | **Protein Consequence** | **Allele Count** | **gnomAD Allele Frequency** | **Primary Source** | **Additional Sources** | **Mean Procko Score** | **Contact Residue** |
| 248 | 706 | M706I | p.Met706Ile | 1 | 7.08572E-06 | gnomAD | | | |
| 249 | 708 | R708Q | p.Arg708Gln | 1 | 6.91247E-06 | gnomAD | Other | | |
| 250 | 708 | R708W | p.Arg708Trp | 3 | 1.80397E-05 | gnomAD | | | |
| 251 | 709 | S709R | p.Ser709Arg | 1 | 6.88108E-06 | gnomAD | | | |
| 252 | 710 | R710C | p.Arg710Cys | 5 | 2.89195E-05 | gnomAD | Other | | |
| 253 | 710 | R710H | p.Arg710His | 7 | 3.99657E-05 | gnomAD | | | |
| 254 | 716 | R716H | p.Arg716His | 15 | 8.18027E-05 | gnomAD | Other | | |
| 255 | 716 | R716C | p.Arg716Cys | 1 | 6.19149E-06 | gnomAD | | | |
| 256 | 719 | D719E | p.Asp719Glu | 1 | 6.01525E-06 | gnomAD | | | |
| 257 | 720 | N720D | p.Asn720Asp | 3054 | 0.016215011 | gnomAD | AsianSpecificDB,UKBio | | |
| 258 | 720 | N720S | p.Asn720Ser | 1 | 6.00402E-06 | gnomAD | | | |
| 259 | 726 | G726R | p.Gly726Arg | 2 | 1.15294E-05 | gnomAD | | | |
| 260 | 726 | G726E | p.Gly726Glu | 1 | 5.76558E-06 | gnomAD | | | |
| 261 | 729 | P729L | p.Pro729Leu | 2 | 1.00837E-05 | gnomAD | | | |
| 262 | 730 | T730K | p.Thr730Lys | 1 | 5.63146E-06 | gnomAD | | | |
| 263 | 731 | L731F | p.Leu731Phe | 286 | 0.001434915 | gnomAD | AsianSpecificDB,UKBio | | |
| 264 | 733 | P733L | p.Pro733Leu | 2 | 0 | AsianSpecif | AsianSpecificDB | | |
| 265 | 734 | P734L | p.Pro734Leu | 2 | 1.12724E-05 | gnomAD | AsianSpecificDB | | |
| 266 | 735 | N735K | p.Asn735Lys | 1 | 5.64127E-06 | gnomAD | | | |
| 267 | 737 | P737A | p.Pro737Ala | 1 | 5.64723E-06 | gnomAD | AsianSpecificDB | | |
| 268 | 737 | P737L | p.Pro737Leu | 2 | 0 | AsianSpecif | AsianSpecificDB | | |
| 269 | 740 | S740P | p.Ser740Pro | 1 | 4.59749E-05 | gnomAD | | | |
| 270 | 741 | I741V | p.Ile741Val | 20 | 0.000100345 | gnomAD | Other | | |
| 271 | 745 | V745I | p.Val745Ile | 2 | 1.01287E-05 | gnomAD | | | |
| 272 | 751 | G751E | p.Gly751Glu | 2 | 1.1659E-05 | gnomAD | | | |
| 273 | 752 | V752M | p.Val752Met | 1 | 0 | AsianSpecif | AsianSpecificDB | | |
| 274 | 753 | I753M | p.Ile753Met | 1 | 5.83216E-06 | gnomAD | AsianSpecificDB | | |
| 275 | 761 | I761V | p.Ile761Val | 1 | 6.28943E-06 | gnomAD | | | |
| 276 | 767 | D767H | p.Asp767His | 2 | 1.34282E-05 | gnomAD | | | |
| 277 | 768 | R768W | p.Arg768Trp | 2 | 1.38522E-05 | gnomAD | | | |
| 278 | 769 | K769E | p.Lys769Glu | 1 | 6.92919E-06 | gnomAD | | | |

| | A | B | C | D | E | F | G | H | I |
|---|---|---|---|---|---|---|---|---|---|
| 2 | **Position** | **Mutation** | **Protein Consequence** | **Allele Count** | **gnomAD Allele Frequency** | **Primary Source** | **Additional Sources** | **Mean Procko Score** | **Contact Residue** |
| 279 | 771 | K771R | p.Lys771Arg | 1 | 4.53762E-05 | gnomAD | | | |
| 280 | 772 | N772S | p.Asn772Ser | 12 | 6.02101E-05 | gnomAD | | | |
| 281 | 774 | A774G | p.Ala774Gly | 1 | 5.61605E-06 | gnomAD | | | |
| 282 | 774 | A774P | p.Ala774Pro | 1 | 5.61549E-06 | gnomAD | | | |
| 283 | 774 | A774T | p.Ala774Thr | 1 | 5.61549E-06 | gnomAD | | | |
| 284 | 776 | S776R | p.Ser776Arg | 1 | 4.55021E-05 | gnomAD | | | |
| 285 | 781 | Y781H | p.Tyr781His | 3 | 1.4797E-05 | gnomAD | | | |
| 286 | 782 | A782V | p.Ala782Val | 11 | 6.07839E-05 | gnomAD | AsianSpecificDB,UKBio | | |
| 287 | 785 | D785N | p.Asp785Asn | 7 | 3.86678E-05 | gnomAD | AsianSpecificDB,UKBio | | |
| 288 | 793 | P793L | p.Pro793Leu | 1 | 5.57324E-06 | gnomAD | | | |
| 289 | 796 | Q796R | p.Gln796Arg | 2 | 1.11324E-05 | gnomAD | | | |
| 290 | 801 | V801G | p.Val801Gly | 1 | 5.61728E-06 | gnomAD | AsianSpecificDB,UKBio | | |
| 291 | 802 | Q802R | p.Gln802Arg | 1 | 0 | AsianSpecif | AsianSpecificDB | | |
| 292 | 804 | S804F | p.Ser804Phe | 1 | 5.80124E-06 | gnomAD | | | |
| 293 | 805 | F805I | p.Phe805Ile | 4 | 2.3209E-05 | gnomAD | | | |
| 294 | 482 | R482* | p.Arg482Ter | 1 | 0 | Other | | | |
| 295 | 244 | V244fs | p.Val244LeufsTe | 1 | 0 | Other | | | |
| 296 | 656 | L656* | p.Leu656Ter | 1 | 5.9153E-06 | gnomAD | AsianSpecificDB | | |
| 297 | 405 | G405del | p.Gly405del | 2 | 1.09215E-05 | gnomAD | | | |
| 298 | 422 | G422fs | p.Gly422ValfsTe | 1 | 5.46472E-06 | gnomAD | | | |
| 299 | 116 | L116* | p.Leu116Ter | 1 | 6.43882E-06 | gnomAD | | | |
| 300 | 313 | K313del | p.Lys313del | 1 | 5.642E-06 | gnomAD | | | |

### Supplementary Table 2 - ACE2 polymorphisms predicted to alter SARS-CoV/Cov-2 S-protein binding

| | A | B | C | D | E | F | G | H | I | J | K | L |
|---|---|---|---|---|---|---|---|---|---|---|---|---|
| 1 | Supplementary Table 2 - ACE2 polymorphisms predicted to alter SARS-CoV/Cov-2 S-protein binding | | | | | | | | | | | |
| 2 | Position | Mutation | Protein Consequence | Allele Count | gnomAD Allele Frequency | Primary Source | Additional Sources | Mean enrichment score (Procko 2020) | Contact Residue | Strctural mechanism (Fig 2) | impact (binding to S-protein) | Suceptibility to SARS-CoV/SRAS-CoV-2 (combined interpretation) |
| 3 | 38 | D38V | p.Asp38Val | 1 | 0 | Other | | -1.9637347 | Y | Direct | Decrease | Decrease |
| 4 | 31 | K31R | p.Lys31Arg | 1 | 0 | AsianSpecifi | AsianSpecifi | -1.6094772 | Y | Direct | Decrease | Decrease |
| 5 | 33 | N33I | p.Asn33Ile | 1 | 0 | AsianSpecifi | AsianSpecifi | -1.5683037 | | Direct | Decrease | Decrease |
| 6 | 37 | E37K | p.Glu37Lys | 8 | 3.897E-05 | gnomAD | AsianSpecifi | -1.3499866 | Y | Indirect | Decrease | Decrease |
| 7 | 35 | E35K | p.Glu35Lys | 3 | 1.636E-05 | gnomAD | | -1.1992532 | Y | Direct | Decrease | Decrease |
| 8 | 388 | Q388L | p.Gln388Leu | 4 | 2.186E-05 | gnomAD | | -1.0844672 | | Indirect | Decrease | Decrease |
| 9 | 34 | H34R | p.His34Arg | 1 | 0 | AsianSpecifi | AsianSpecifi | -1.0760313 | Y | Direct | Decrease | Decrease |
| 10 | 51 | N51S | p.Asn51Ser | 1 | 5.488E-06 | gnomAD | | -1.0527784 | | Indirect | Decrease | Decrease |
| 11 | 68 | K68E | p.Lys68Glu | 2 | 1.095E-05 | gnomAD | AsianSpecifi | -1.052667 | | Indirect | Decrease | Decrease |
| 12 | 352 | G352V | p.Gly352Val | 1 | 5.752E-06 | gnomAD | | -1.0253194 | | Direct | Decrease | Decrease |
| 13 | 509 | D509Y | p.Asp509Tyr | 1 | 0 | Other | | -0.9429219 | | - | - | - |
| 14 | 355 | D355N | p.Asp355Asn | 2 | 1.174E-05 | gnomAD | Other | -0.9110524 | | Indirect | Decrease | Decrease |
| 15 | 326 | G326E | p.Gly326Glu | 1 | 5.517E-06 | gnomAD | | -0.9054689 | Y | Direct | Decrease | Decrease |
| 16 | 72 | F72V | p.Phe72Val | 1 | 5.474E-06 | gnomAD | | -0.8867353 | | Indirect | Decrease | Decrease |
| 17 | 62 | M62V | p.Met62Val | 1 | 5.666E-06 | gnomAD | AsianSpecifi | -0.7827955 | | Indirect | Decrease | Decrease |
| 18 | 83 | Y83H | p.Tyr83His | 2 | 0 | SAS(1) | AsianSpecifi | -0.6671349 | Y | Direct | Decrease | Decrease |
| 19 | 50 | Y50F | p.Tyr50Phe | 1 | 5.482E-06 | gnomAD | | -0.5353604 | | Indirect | Decrease | Decrease |
| 20 | 329 | E329G | p.Glu329Gly | 7 | 3.443E-05 | gnomAD | Other | -0.5064661 | Y | - | Decrease | Decrease |
| 21 | 51 | N51D | p.Asn51Asp | 1 | 5.485E-06 | gnomAD | | -0.5008654 | | - | Decrease | Decrease |
| 22 | 62 | M62I | p.Met62Ile | 1 | 0 | AsianSpecifi | AsianSpecifi | -0.4332215 | | - | Decrease | Decrease |
| 23 | 82 | M82I | p.Met82Ile | 5 | 2.442E-05 | gnomAD | | -0.4285018 | Y | - | Decrease | Decrease |
| 24 | 84 | P84T | p.Pro84Thr | 1 | 5.471E-06 | gnomAD | | -0.4191317 | | - | Decrease | Decrease |
| 25 | 389 | P389H | p.Pro389His | 7 | 3.826E-05 | gnomAD | | -0.4007303 | | - | Decrease | Decrease |
| 26 | 290 | N290H | p.Asn290His | 2 | 1.133E-05 | gnomAD | | -0.3820868 | | - | - | - |
| 27 | 518 | R518T | p.Arg518Thr | 1 | 5.49E-06 | gnomAD | | -0.3408209 | | - | - | - |
| 28 | 504 | F504I | p.Phe504Ile | 2 | 1.268E-05 | gnomAD | | -0.3318035 | | - | - | - |
| 29 | 511 | S511P | p.Ser511Pro | 1 | 0 | AsianSpecifi | AsianSpecifi | -0.0965233 | | - | - | - |
| 30 | 346 | P346S | p.Pro346Ser | 1 | 5.599E-06 | gnomAD | | -0.0669063 | | - | - | - |
| 31 | 504 | F504L | p.Phe504Leu | 1 | 4.631E-05 | gnomAD | | -0.0355608 | | - | - | - |
| 32 | 366 | M366T | p.Met366Thr | 2 | 1.098E-05 | gnomAD | | 0.0341968 | | - | - | - |

| | A | B | C | D | E | F | G | H | I | J | K | L |
|---|---|---|---|---|---|---|---|---|---|---|---|---|
| 2 | **Position** | **Mutation** | **Protein Consequence** | **Allele Count** | **gnomAD Allele Frequency** | **Primary Source** | **Additional Sources** | **Mean enrichment score (Procko 2020)** | **Contact Residue** | **Strctural mechanism (Fig 2)** | **impact (binding to S-protein)** | **Suceptibility to SARS-CoV/SRAS-CoV-2 (combined interpretation )** |
| 33 | 446 | I446M | p.Ile446Met | 1 | 4.54E-05 | gnomAD | | 0.0591718 | | - | - | - |
| 34 | 21 | I21T | p.Ile21Thr | 1 | 5.463E-06 | gnomAD | | 0.072108 | | - | - | - |
| 35 | 374 | H374R | p.His374Arg | 1 | 5.475E-06 | gnomAD | | 0.1107117 | | - | - | - |
| 36 | 398 | E398K | p.Glu398Lys | 1 | 5.462E-06 | gnomAD | | 0.2057975 | | - | - | - |
| 37 | 510 | Y510H | p.Tyr510His | 1 | 6.863E-06 | gnomAD | | 0.2460174 | | - | - | - |
| 38 | 445 | T445M | p.Thr445Met | 1 | 5.869E-06 | gnomAD | | 0.4226164 | | - | - | - |
| 39 | 40 | F40L | p.Phe40Leu | 2 | 3.086E-05 | Gnomad | | 0.4490517 | | - | Increase | Increased |
| 40 | 26 | K26E | p.Lys26Glu | 1 | 5.455E-06 | gnomAD | | 0.5734929 | | - | Increase | Increased |
| 41 | 60 | Q60R | p.Gln60Arg | 2 | 1.125E-05 | gnomAD | | 0.6028484 | | - | Increase | Increased |
| 42 | 23 | E23K | p.Glu23Lys | 1 | 5.458E-06 | gnomAD | | 0.647777 | | Indirect | Increase | Increased |
| 43 | 383 | M383T | p.Met383Thr | 1 | 0 | AsianSpecifi | AsianSpecifi | 0.6762501 | | - | Increase | Increased |
| 44 | 21 | I21V | p.Ile21Val | 2 | 1.093E-05 | gnomAD | Other | 0.6970124 | | Direct | Increase | Increased |
| 45 | 378 | H378R | p.His378Arg | 18 | 8.791E-05 | gnomAD | Other | 0.8315449 | | - | Increase | Increased |
| 46 | 64 | N64K | p.Asn64Lys | 3 | 1.466E-05 | gnomAD | | 1.0472096 | | - | Increase | Increased |
| 47 | 102 | Q102P | p.Gln102Pro | 3 | 1.475E-05 | gnomAD | | 1.1913283 | | - | Increase | Increased |
| 48 | 26 | K26R | p.Lys26Arg | 797 | 0.0038833 | gnomAD | AsianSpecifi | 1.2292467 | | Direct | Increase | Increased |
| 49 | 27 | T27A | p.Thr27Ala | 2 | 1.091E-05 | gnomAD | | 2.1134249 | Y | Indirect | Increase | Increased |
| 50 | 19 | S19P | p.Ser19Pro | 64 | 0.0003129 | gnomAD | Other | 2.3278184 | | Direct | Increase | Increased |
| 51 | 92 | T92I | p.Thr92Ile | 2 | 1.096E-05 | gnomAD | | 2.5342868 | | Direct | Increase | Increased |

| | A | B | C | D | E | F | G | H |
|---|---|---|---|---|---|---|---|---|
| 1 | **Supplementary Table 3 - Conservation of N90 glycosylation motif in annotated jawed vertebrate ACE2 orthologs** | | | | | | | |
| 2 | | | | | | | | |
| 3 | **Common name** | **Refseq ID** | **Residue #90** | **Residue #90** | **Residue #91** | **Residue #92** | **Residue #93** | **Motif NxT/S present?** |
| 4 | Human | NP_001358344.1 | Q | N | L | T | V | Yes |
| 5 | house mouse | NP_081562.2 | Q | T | P | I | I | No |
| 6 | Norway rat | NP_001012006.1 | Q | N | A | T | I | No |
| 7 | zebrafish | XP_005169416.1 | S | D | P | I | I | No |
| 8 | pig | NP_001116542.1 | Q | T | L | I | L | No |
| 9 | Rhesus monke | NP_001129168.1 | Q | N | L | T | V | Yes |
| 10 | cattle | XP_005228485.1 | Q | N | L | T | L | Yes |
| 11 | dog | NP_001158732.1 | Q | D | S | T | V | No |
| 12 | rabbit | XP_002719891.1 | Q | N | L | T | V | Yes |
| 13 | tropical clawed | XP_002938293.2 | T | D | P | S | I | No |
| 14 | chicken | XP_416822.2 | Q | D | A | V | T | No |
| 15 | chimpanzee | XP_016798468.1 | Q | N | L | T | V | Yes |
| 16 | domestic cat | XP_023104564.1 | H | N | T | T | V | Yes |
| 17 | sheep | XP_011961657.1 | Q | N | L | T | L | Yes |
| 18 | rainbow trout | XP_021433278.1 | S | D | P | L | I | No |
| 19 | Atlantic cod | XP_030232530.1 | K | D | P | V | V | No |
| 20 | giant panda | XP_002930657.1 | H | N | S | T | V | Yes |
| 21 | Brandt's bat | XP_014399780.1 | Q | N | L | T | I | Yes |
| 22 | elephant shark | XP_007889845.1 | S | D | N | I | I | No |
| 23 | domestic ferret | NP_001297119.1 | Q | D | P | I | I | No |
| 24 | golden hamste | XP_005074266.1 | Q | N | L | T | I | Yes |
| 25 | naked mole-rat | XP_004866157.1 | Q | N | L | T | V | Yes |
| 26 | wild yak | XP_005903173.1 | Q | N | L | T | L | Yes |
| 27 | barramundi per | XP_018539189.1 | K | D | Q | E | I | No |
| 28 | white-tufted-ea | XP_008987241.1 | Q | N | L | T | V | Yes |
| 29 | horse | XP_001490241.1 | Q | N | L | T | V | Yes |
| 30 | greater amberj | XP_022605054.1 | K | N | P | E | I | No |

| | A | B | C | D | E | F | G | H |
|---|---|---|---|---|---|---|---|---|
| 3 | **Common name** | **Refseq ID** | **Residue #90** | **Residue #90** | **Residue #91** | **Residue #92** | **Residue #93** | **Motif NxT/S present?** |
| 31 | turquoise killifis | XP_015808977.1 | K | D | P | E | V | No |
| 32 | two-lined caecili | XP_029459086.1 | T | E | P | E | I | No |
| 33 | Microcaecilia un | XP_030058174.1 | T | D | P | E | T | No |
| 34 | Mexican tetra | XP_022523929.1 | S | D | E | L | V | No |
| 35 | coelacanth | XP_005997915.2 | T | D | P | H | I | No |
| 36 | spotted gar | XP_006639185.1 | A | D | K | K | I | No |
| 37 | Atlantic herring | XP_031414786.1 | N | D | L | E | I | No |
| 38 | goldfish | XP_026131313.1 | S | D | P | L | I | No |
| 39 | channel catfish | XP_017313836.1 | S | D | H | E | V | No |
| 40 | electric eel | XP_026867211.1 | T | D | P | E | I | No |
| 41 | northern pike | XP_010884777.1 | K | D | P | L | I | No |
| 42 | Atlantic salmon | XP_014062928.1 | . | . | . | . | . | NA |
| 43 | Arctic char | XP_023998967.1 | S | V | I | I | D | No |
| 44 | mummichog | XP_021178197.1 | K | D | P | Q | I | No |
| 45 | guppy | XP_008402714.1 | S | D | P | V | I | No |
| 46 | southern platyfi | XP_005799835.1 | N | D | P | V | I | No |
| 47 | Nile tilapia | XP_003445853.2 | N | D | L | E | I | No |
| 48 | Burton's mouth | XP_005943362.1 | N | D | L | E | I | No |
| 49 | eastern happy | XP_026020155.1 | N | D | L | E | I | No |
| 50 | yellow perch | XP_028441363.1 | K | D | P | E | I | No |
| 51 | gilthead seabre | XP_030271236.1 | K | D | R | E | L | No |
| 52 | black rockcod | XP_010790455.1 | T | D | A | T | I | No |
| 53 | Japanese floun | XP_019935235.1 | K | D | A | K | I | No |
| 54 | Green sea turtl | XP_007070561.1 | M | D | P | I | V | No |
| 55 | Painted turtle | XP_023964517.1 | T | D | P | I | V | No |
| 56 | American alliga | XP_019350687.1 | M | D | P | L | I | No |
| 57 | Australian saltw | XP_019384826.1 | . | D | P | V | I | No |
| 58 | mainland tiger | XP_026530754.1 | S | N | E | T | I | Yes |
| 59 | Pseudonaja tex | XP_026570054.1 | A | N | E | T | I | Yes |
| 60 | emu | XP_025976569.1 | T | D | D | L | I | No |
| 61 | mallard | XP_012949915.2 | Q | D | P | L | L | No |
| 62 | chimney swift | XP_009992128.1 | S | D | A | L | I | No |

| | A | B | C | D | E | F | G | H |
|---|---|---|---|---|---|---|---|---|
| 3 | **Common name** | **Refseq ID** | **Residue #90** | **Residue #90** | **Residue #91** | **Residue #92** | **Residue #93** | **Motif NxT/S present?** |
| 63 | rock pigeon | XP_021154486.1 | Q | D | D | L | T | No |
| 64 | peregrine falco | XP_005231984.2 | Q | D | A | L | T | No |
| 65 | white-tailed eag | XP_009925641.1 | Q | D | D | L | T | No |
| 66 | helmeted guine | XP_021240731.1 | Q | D | A | V | T | No |
| 67 | Ring-necked ph | XP_031451919.1 | Q | D | A | A | T | No |
| 68 | turkey | XP_019467554.1 | Q | D | A | A | T | No |
| 69 | Common canar | XP_009087922.1 | K | D | D | L | T | No |
| 70 | Great Tit | XP_015486815.1 | T | D | D | L | T | No |
| 71 | Common starlir | XP_014731370.1 | T | D | D | L | I | No |
| 72 | great cormoran | XP_009509070.1 | Q | D | A | L | T | No |
| 73 | emperor pengu | XP_009275140.1 | Q | D | A | L | T | No |
| 74 | Adelie penguin | XP_009323767.1 | Q | D | T | L | T | No |
| 75 | Anna's hummir | XP_008492997.2 | T | D | A | L | I | No |
| 76 | platypus | XP_001515597.2 | S | D | R | S | L | No |
| 77 | Tasmanian dev | XP_031814825.1 | S | A | Y | P | I | No |
| 78 | nine-banded ar | XP_004449124.1 | S | N | L | T | N | Yes |
| 79 | western Europe | XP_007538670.1 | Q | N | P | T | V | No |
| 80 | small Madagasca | XP_004710002.1 | T | D | P | I | I | No |
| 81 | black flying fox | XP_006911709.1 | Q | D | P | I | L | No |
| 82 | Egyptian rouse | XP_015974412.1 | Q | D | P | E | L | No |
| 83 | common vampi | XP_024425698.1 | K | D | V | N | V | No |
| 84 | Tufted capuchi | XP_032141854.1 | Q | N | L | T | V | Yes |
| 85 | sooty mangabe | XP_011891198.1 | Q | N | L | T | V | Yes |
| 86 | crab-eating ma | XP_005593094.1 | Q | N | L | T | V | Yes |
| 87 | pig-tailed maca | XP_011733505.1 | Q | N | L | T | V | Yes |
| 88 | olive baboon | XP_021788732.1 | Q | N | L | T | V | Yes |
| 89 | gelada | XP_025227847.1 | Q | N | L | T | V | Yes |
| 90 | drill | XP_011850923.1 | Q | N | L | T | V | Yes |
| 91 | western gorilla | XP_018874749.1 | Q | N | L | T | I | Yes |
| 92 | pygmy chimpar | XP_008972428.1 | Q | N | L | T | V | Yes |
| 93 | Sumatran oran | NP_001124604.1 | Q | N | L | T | V | Yes |
| 94 | red fox | XP_025842512.1 | Q | D | S | T | V | No |

| | A | B | C | D | E | F | G | H |
|---|---|---|---|---|---|---|---|---|
| 3 | **Common name** | **Refseq ID** | **Residue #90** | **Residue #90** | **Residue #91** | **Residue #92** | **Residue #93** | **Motif NxT/S present?** |
| 95 | leopard | XP_019273508.1 | H | N | T | T | V | Yes |
| 96 | puma | XP_025790417.1 | H | N | T | T | V | Yes |
| 97 | California sea lio | XP_027465353.1 | Q | D | S | T | V | No |
| 98 | Pacific walrus | XP_004415448.1 | Q | D | S | T | V | No |
| 99 | Weddell seal | XP_030886750.1 | . | . | . | . | . | NA |
| 100 | harbor seal | XP_032245506.1 | Q | D | S | T | V | No |
| 101 | long-finned pilo | XP_030703991.1 | R | N | L | T | L | Yes |
| 102 | killer whale | XP_004269705.1 | R | N | L | T | L | Yes |
| 103 | common bottle | XP_019781177.1 | R | N | L | T | L | Yes |
| 104 | beluga whale | XP_022418360.1 | R | N | L | T | L | Yes |
| 105 | sperm whale | XP_023971279.1 | Q | N | L | T | L | Yes |
| 106 | African savann | XP_023410960.1 | S | S | S | I | I | No |
| 107 | ass | XP_014713133.1 | Q | N | L | T | V | Yes |
| 108 | Przewalski's ho | XP_008542995.1 | Q | N | L | T | V | Yes |
| 109 | Bactrian camel | XP_010966303.1 | Q | N | V | T | L | Yes |
| 110 | Arabian camel | XP_010991717.1 | Q | N | V | T | L | Yes |
| 111 | Odocoileus virg | XP_020768965.1 | Q | N | L | T | L | Yes |
| 112 | zebu cattle | XP_019811719.1 | Q | N | L | T | L | Yes |
| 113 | goat | NP_001277036.1 | Q | N | L | T | L | Yes |
| 114 | Malayan pango | XP_017505746.1 | Q | N | D | T | I | Yes |
| 115 | American pika | XP_004597549.2 | Q | N | L | T | T | Yes |
| 116 | Alpine marmot | XP_015343540.1 | Q | N | F | T | L | Yes |
| 117 | Arctic ground s | XP_026252505.1 | Q | N | F | T | L | Yes |
| 118 | Ord's kangaroo | XP_012887572.1 | Q | N | P | I | L | No |
| 119 | Chinese hamst | XP_003503283.1 | Q | N | L | I | I | No |
| 120 | white-footed m | XP_028743609.1 | P | N | L | I | I | No |
| 121 | Ryukyu mouse | XP_021009138.1 | Q | T | P | I | I | No |
| 122 | shrew mouse | XP_021043935.1 | Q | N | P | V | I | No |
| 123 | domestic guine | XP_023417808.1 | Q | N | L | T | V | Yes |
| 124 | degu | XP_023575315.1 | Q | N | L | T | V | Yes |
| 125 | gray short-taile | XP_007500935.1 | T | N | A | T | V | Yes |
| 126 | Chinese soft-sh | XP_006122891.1 | T | N | H | T | V | Yes |

| | A | B | C | D | E | F | G | H |
|---|---|---|---|---|---|---|---|---|
| 3 | **Common name** | **Refseq ID** | **Residue #90** | **Residue #90** | **Residue #91** | **Residue #92** | **Residue #93** | **Motif NxT/S present?** |
| 127 | Bolivian squirrel | XP_010334925.1 | Q | N | L | T | V | Yes |
| 128 | reedfish | XP_028655640.1 | S | N | Y | T | I | Yes |
| 129 | green anole | XP_008105455.1 | N | N | D | T | I | Yes |
| 130 | Cape elephant | XP_006892457.1 | S | D | P | S | I | No |
| 131 | sheepshead mi | XP_015226730.1 | K | D | L | Q | I | No |
| 132 | polar bear | XP_008694637.1 | H | N | S | T | V | Yes |
| 133 | big brown bat | XP_008153150.1 | Q | N | L | T | I | Yes |
| 134 | Hawaiian monk | XP_021536480.1 | Q | D | S | T | L | No |
| 135 | common womb | XP_027691156.1 | S | D | P | Q | I | No |
| 136 | Opisthocomus | XP_009938970.1 | Q | D | A | L | T | No |
| 137 | Northern fulma | XP_009574896.1 | H | D | A | L | T | No |
| 138 | Nothoprocta pe | XP_025891105.1 | K | N | D | L | I | No |
| 139 | hybrid cattle | XP_027389727.1 | Q | N | L | T | L | Yes |
| 140 | alpaca | XP_006212709.1 | E | N | V | T | L | Yes |
| 141 | gray mouse len | XP_020140826.1 | Q | N | L | T | I | Yes |
| 142 | small-eared gala | XP_003791912.1 | Q | N | R | T | V | Yes |
| 143 | Indian medaka | XP_024150631.1 | K | D | P | E | I | No |
| 144 | torafugu | XP_029702274.1 | K | N | A | E | I | No |
| 145 | Monterrey platy | XP_027871671.1 | N | D | P | V | I | No |
| 146 | cheetah | XP_026910297.1 | H | N | T | T | V | Yes |
| 147 | long-tailed chin | XP_013362428.1 | Q | N | L | T | V | Yes |
| 148 | northern fur sea | XP_025713397.1 | Q | D | S | T | V | No |
| 149 | Steller sea lion | XP_027970822.1 | Q | D | S | T | V | No |
| 150 | Western terrest | XP_032082934.1 | T | N | E | T | I | Yes |
| 151 | Thamnophis si | XP_013926936.1 | T | N | E | T | I | Yes |
| 152 | southern multin | XP_031226742.1 | K | T | P | I | I | No |
| 153 | Dalmatian pelic | XP_009478920.1 | Q | D | D | L | T | No |
| 154 | ermine | XP_032187677.1 | Q | D | P | I | I | No |
| 155 | mangrove rivulu | XP_017295385.1 | H | D | P | T | V | No |
| 156 | meerkat | XP_029786256.1 | Q | N | T | T | V | Yes |
| 157 | red-throated lo | XP_009816127.1 | Q | D | A | L | I | Yes |
| 158 | Ma's night mon | XP_012290105.1 | Q | N | L | T | V | Yes |

| | A | B | C | D | E | F | G | H |
|---|---|---|---|---|---|---|---|---|
| 3 | **Common name** | **Refseq ID** | **Residue #90** | **Residue #90** | **Residue #91** | **Residue #92** | **Residue #93** | **Motif NxT/S present?** |
| 159 | koala | XP_020863153.1 | S | D | P | Q | I | No |
| 160 | Chinese alligat | XP_025066628.1 | . | D | P | L | I | No |
| 161 | narwhal | XP_029095804.1 | R | N | L | T | L | Yes |
| 162 | European shre | XP_004612266.1 | T | D | P | K | V | No |
| 163 | red-bellied pira | XP_017550079.1 | S | D | P | L | I | No |
| 164 | thirteen-lined g | XP_005316051.3 | Q | N | F | T | L | Yes |
| 165 | Bison bison bis | XP_010833001.1 | Q | N | L | T | L | Yes |
| 166 | swamp eel | XP_020465646.1 | R | D | A | E | I | No |
| 167 | white-throated | XP_005491832.2 | T | D | E | L | T | No |
| 168 | Oreochromis a | XP_031584810.1 | N | D | L | E | I | No |
| 169 | Amazon molly | XP_007560208.1 | N | D | P | V | I | No |
| 170 | sailfin molly | XP_014895313.1 | N | D | P | V | I | No |
| 171 | Poecilia mexica | XP_014837025.1 | N | D | P | V | I | No |
| 172 | medium ground | XP_005426221.1 | T | D | E | L | T | No |
| 173 | killdeer | XP_009887331.1 | Q | D | P | L | I | No |
| 174 | lesser Egyptian | XP_004671523.1 | Q | N | P | T | I | No |
| 175 | bald eagle | XP_010579828.1 | Q | D | D | L | T | No |
| 176 | Austrofundulus | XP_013888928.1 | N | D | P | N | I | No |
| 177 | brown roatelo | XP_010178703.1 | Q | D | P | L | I | No |
| 178 | Red-legged se | XP_009703695.1 | Q | D | A | L | T | No |
| 179 | sunbittern | XP_010156467.1 | Q | D | A | L | I | No |
| 180 | common cucko | XP_009563864.1 | Q | D | A | L | T | No |
| 181 | Barn owl | XP_009969209.1 | Q | D | A | L | T | No |
| 182 | Cottoperca gob | XP_029283581.1 | R | D | S | T | I | No |
| 183 | ballan wrasse | XP_020493627.1 | T | I | P | E | I | No |
| 184 | rifleman | XP_009082150.1 | . | . | . | . | . | NA |
| 185 | bar-tailed trogo | XP_009867056.1 | G | D | D | L | I | No |
| 186 | speckled mous | XP_010206054.1 | . | . | . | . | . | NA |
| 187 | carmine bee-ea | XP_008937519.1 | Q | N | A | T | T | Yes |
| 188 | little brown bat | XP_023609437.1 | Q | N | S | T | I | Yes |
| 189 | zebra finch | XP_002194303.3 | A | D | D | P | T | No |
| 190 | collared flycatc | XP_005037422.1 | T | D | D | L | T | No |

| | A | B | C | D | E | F | G | H |
|---|---|---|---|---|---|---|---|---|
| 3 | Common name | Refseq ID | Residue #90 | Residue #90 | Residue #91 | Residue #92 | Residue #93 | Motif NxT/S present? |
| 191 | green monkey | XP_007989304.1 | Q | N | L | T | V | Yes |
| 192 | Canada lynx | XP_030160839.1 | H | N | T | T | I | Yes |
| 193 | black snub-nos | XP_017744069.1 | Q | N | L | T | V | Yes |
| 194 | golden snub-nc | XP_010364367.2 | Q | N | L | T | V | Yes |
| 195 | | XP_003261132.2 | Q | N | L | T | I | Yes |
| 196 | flier cichlid | XP_030582139.1 | Q | D | L | E | I | No |
| 197 | climbing perch | XP_026233431.1 | S | D | P | E | I | No |
| 198 | Amur tiger | XP_007090142.1 | H | N | T | T | V | Yes |
| 199 | prairie vole | XP_005358818.1 | Q | N | L | L | L | No |
| 200 | spiny chromis | XP_022063988.1 | T | D | P | E | I | No |
| 201 | clown anemone | XP_023124156.1 | K | D | P | E | I | No |
| 202 | American crow | XP_017583883.1 | . | . | . | . | . | NA |
| 203 | Camarhynchus | XP_030811385.1 | T | D | E | L | T | No |
| 204 | water buffalo | XP_006041602.1 | Q | N | L | T | L | Yes |
| 205 | pale spear-nos | XP_028378317.1 | T | D | V | T | V | No |
| 206 | Pacific white-si | XP_026951598.1 | R | N | L | T | L | Yes |
| 207 | yellow-bellied r | XP_027802308.1 | Q | N | F | T | L | Yes |
| 208 | Japanese quail | XP_015742063.1 | Q | D | A | V | T | No |
| 209 | white-throated | XP_010217584.1 | K | D | D | L | I | No |
| 210 | Gharial | XP_019381060.1 | . | D | P | L | I | No |
| 211 | White-tailed tro | XP_010290019.1 | Q | D | A | L | T | No |
| 212 | central bearded | XP_020642422.1 | S | N | E | T | I | Yes |
| 213 | Protobothrops | XP_029140508.1 | T | N | E | T | I | Yes |
| 214 | zebra mbuna | XP_004543482.1 | N | D | L | E | I | No |
| 215 | tiger tail seahor | XP_019742561.1 | K | D | P | Q | I | No |
| 216 | Asian bonytong | XP_018584732.1 | T | D | P | T | V | No |
| 217 | saffron-crested | XP_027544864.1 | E | D | N | L | I | No |
| 218 | Ursus arctos ho | XP_026333865.1 | H | N | S | T | V | Yes |
| 219 | Downy woodpe | XP_009909849.1 | . | . | . | . | . | NA |
| 220 | Yangtze River | XP_007466389.1 | Q | N | L | T | L | Yes |
| 221 | red-crested tura | XP_009978415.1 | Q | D | A | L | I | No |
| 222 | Nanorana park | XP_018418558.1 | T | D | E | M | L | No |

| | A | B | C | D | E | F | G | H |
|---|---|---|---|---|---|---|---|---|
| 3 | **Common name** | **Refseq ID** | **Residue #90** | **Residue #90** | **Residue #91** | **Residue #92** | **Residue #93** | **Motif NxT/S present?** |
| 223 | crested ibis | XP_009474590.1 | Q | D | A | L | T | No |
| 224 | large flying fox | XP_011361275.1 | Q | D | P | I | L | No |
| 225 | star-nosed mol | XP_012585871.1 | Q | D | P | I | V | No |
| 226 | bicolor damself | XP_008290762.1 | K | D | P | E | I | No |
| 227 | Gekko japonicu | XP_015273067.1 | S | D | P | H | I | No |
| 228 | great blue-spot | XP_020781598.1 | K | D | R | E | I | No |
| 229 | blue tit | XP_023774184.1 | T | D | D | L | T | No |
| 230 | Siamese fightin | XP_028999570.1 | S | D | P | E | I | No |
| 231 | willow flycatche | XP_027757151.1 | E | D | N | L | I | No |
| 232 | live sharksucke | XP_029354066.1 | K | D | P | E | I | No |
| 233 | Buceros rhinoc | XP_010136813.1 | Q | H | D | L | T | No |
| 234 | Kea | XP_010012481.1 | . | . | . | . | . | NA |
| 235 | Burmese pytho | XP_007431942.2 | T | D | E | T | I | No |
| 236 | Tibetan ground | XP_005516712.1 | T | D | D | L | T | No |
| 237 | jewelled blenny | XP_029949252.1 | S | D | P | E | I | No |
| 238 | Cape golden m | XP_006835673.1 | S | N | S | T | I | Yes |
| 239 | great roundleaf | XP_019522936.1 | Q | N | A | T | I | Yes |
| 240 | Macqueen's bu | XP_010120523.1 | Q | D | A | L | T | No |
| 241 | cuckoo roller | XP_009954393.1 | Q | D | A | L | T | No |
| 242 | little egret | XP_009638257.1 | E | D | D | A | T | No |
| 243 | burrowing owl | XP_026705725.1 | Q | D | A | L | T | No |
| 244 | ruff | XP_014815705.1 | Q | D | D | L | T | No |
| 245 | Apteryx mantel | XP_013805736.1 | K | D | D | L | I | No |
| 246 | zig-zag eel | XP_026175949.1 | R | D | P | E | I | No |
| 247 | Indian glassy fi | XP_028257887.1 | T | D | P | E | I | No |
| 248 | large yellow cro | XP_010730146.1 | K | N | P | I | I | No |
| 249 | Aquila chrysae | XP_011587755.1 | Q | D | D | L | T | No |
| 250 | tufted duck | XP_032058386.1 | Q | D | P | L | L | No |
| 251 | Aquila chrysae | XP_029855025.1 | Q | D | D | L | T | No |
| 252 | Myotis davidii | XP_006775273.1 | Q | N | P | T | I | Yes |
| 253 | prairie deer mo | XP_006973269.1 | Q | N | L | I | I | No |
| 254 | yellow-throated | XP_010084373.1 | Q | D | V | L | T | No |

| | A | B | C | D | E | F | G | H |
|---|---|---|---|---|---|---|---|---|
| 3 | **Common name** | **Refseq ID** | **Residue #90** | **Residue #90** | **Residue #91** | **Residue #92** | **Residue #93** | **Motif NxT/S present?** |
| 255 | tongue sole | XP_016887914.1 | K | D | P | E | I | No |
| 256 | Chinese tree sh | XP_006164754.1 | Q | D | T | T | E | No |
| 257 | chuck-will's-wid | XP_010169238.1 | Q | D | A | L | I | No |
| 258 | pike-perch | XP_031162227.1 | K | D | P | E | I | No |
| 259 | dingo | XP_025292925.1 | Q | D | S | T | V | No |
| 260 | Miniopterus nat | XP_016058453.1 | Q | N | S | S | T | Yes |
| 261 | Bengalese finch | XP_021388026.1 | A | D | D | P | T | No |
| 262 | denticle herring | XP_028837781.1 | T | D | P | T | N | No |
| 263 | Cichlid | XP_005724169.1 | N | D | L | E | I | No |
| 264 | Okarito brown k | XP_025942946.1 | K | D | D | L | I | No |
| 265 | Balaenoptera a | XP_028020351.1 | Q | N | L | T | L | Yes |
| 266 | striped catfish | XP_026803610.1 | S | D | Q | E | I | No |
| 267 | blue-crowned m | XP_017667729.1 | E | D | N | L | I | No |
| 268 | golden-collared | XP_017939494.2 | D | D | N | L | I | No |
| 269 | Saker falcon | XP_005443093.2 | Q | D | A | L | T | No |
| 270 | Coquerel's sifa | XP_012494185.1 | Q | N | V | T | V | Yes |
| 271 | Anser cygnoide | XP_013039300.1 | Q | D | P | L | I | No |
| 272 | White-ruffed ma | XP_027494818.1 | E | D | N | L | I | No |
| 273 | Wild Bactrian c | XP_006194263.1 | Q | N | V | T | L | Yes |
| 274 | wolf-eel | XP_031702716.1 | N | D | T | K | I | No |
| 275 | blunt-snouted c | XP_028297875.1 | T | D | L | G | I | No |
| 276 | Struthio camelu | XP_009667495.1 | N | D | D | L | I | No |
| 277 | Grammomys su | XP_028617961.1 | K | T | P | I | I | No |
| 278 | pinecone soldie | XP_029904152.1 | T | D | P | E | I | No |
| 279 | Ugandan red C | XP_023054821.1 | Q | N | L | T | V | Yes |
| 280 | Wire-tailed man | XP_027593974.1 | D | D | N | L | I | No |
| 281 | Damara mole-r | XP_010643477.1 | Q | N | L | T | V | Yes |
| 282 | Corvus cornix c | XP_010392735.2 | T | D | D | L | T | No |
| 283 | Upper Galilee r | XP_008839098.1 | Q | D | L | V | I | No |
| 284 | New Caledonia | XP_031956594.1 | T | D | D | L | T | No |
| 285 | Orycteropus af | XP_007951028.1 | S | N | S | T | I | Yes |
| 286 | yellow catfish | XP_027024524.1 | S | D | P | E | T | No |

| | A | B | C | D | E | F | G | H |
|---|---|---|---|---|---|---|---|---|
| 3 | **Common name** | **Refseq ID** | **Residue #90** | **Residue #90** | **Residue #91** | **Residue #92** | **Residue #93** | **Motif NxT/S present?** |
| 287 | Sinocyclocheilu | XP_016345325.1 | S | D | P | L | I | No |
| 288 | Paramormyrop | XP_023679669.1 | T | D | P | T | I | No |
| 289 | Yangtze finless | XP_024599894.1 | R | N | P | T | L | No |
| 290 | Cebus capucin | XP_017367865.1 | Q | N | L | T | V | Yes |
| 291 | Goodes thorns | XP_030407881.1 | T | D | P | I | V | No |
| 292 | Seriola lalandi ( | XP_023257445.1 | K | N | P | E | I | No |
| 293 | Philippine tarsi | XP_008062810.1 | Q | N | S | T | I | Yes |
| 294 | Kakapo | XP_030332639.1 | E | D | A | L | T | No |
| 295 | budgerigar | XP_005151516.1 | Q | D | A | L | T | No |
| 296 | southern white | XP_004435206.1 | Q | N | V | T | V | Yes |
| 297 | Florida manate | XP_004386381.1 | S | S | S | V | I | No |
| 298 | Enhydra lutris k | XP_022374078.1 | Q | D | P | I | N | No |

### Supplementary Table 4 - Structure evaluation findings

| Mutation | Categorization | Allele Count | Rationale |
|---|---|---|---|
| S19P | Direct Enhancing | 64 | In PDB 6LZG, S19 forms backbone interactions with RBD A475 and backbone interactions with E25 of the same helix. Although these interactions may be stabilizing, prolines are well suited for N-terminal capping of helices and the energetic cost of binding could outstrip the moderate gain achieved through one polar contact. By maintaining more of the helix, S19P could remove this slight steric hindrance where the RBD ridge protrudes. |
| I21V | Indirect Enhancing | 2 | Likely weakens hydrophobic interaction with ACE2 P84 and reduces steric constraints for the bulky ACE2 Y83 to establish hydrophobic interactions with RBD F486 and polar contact with RBD N487. May also facilitate S19 interaction with the RBD. |
| T27A | Direct Enhancing | 2 | T27A removes side-backbone and backbone-backbone interactions. This could endow a1 with greater flexibility to accommodate the RBD ridge binding loop. |
| K26R | Direct Enhancing | 797 | K26R destabilizes N90 glycan while stabilizing intrahelical interaction with ACE2 D30. The N90 glycan may sterically hinder viral recognition. |
| T92I | Direct Enhancing | 2 | Prevents N90 glycosylation by mutating NxS/T glycosylation motif. Additionally, T92 is a helix capping residue. T92I would likely destabilize helix and presentation of the N90-linked glycan. |
| E23K | Indirect Enhancing | 1 | E23 is important for helical fold and likely positions K26 away for interaction with the N90 glycan. |
| M383T | Indirect Enhancing | 1 | M383T appears to stabilize the overall fold with sidechain polar contacts with I379 and Q380. |
| K31R | Direct disrupting | 1 | Loss of polar contact with CoV2 RBD Q493. This residue is distinct from CoV and represents a strengthened interaction. |
| N33I | Indirect disrupting | 1 | N33 forms backbone-backbone and sidechain-backbone contacts with ACE2 L29. Coordinates polar contacts with an $H_2$0 with ACE2 Q96 which helps to appropriately position N90 containing helix. Loss of these interactions could destabilize the central concave portion of ACE2 a1 helix resulting in loss of RBD affinity. |
| H34R | Direct disrupting | 1 | H34 establishes a polar contact with RBD Y453 and intramolecularly contacts D30. H34R would abolish both of those interactions critical for affinity. |
| E35K | Direct disrupting | 3 | E35K likely results in loss of polar contact with RBD Q493. |

| | A | B | C | D |
|---|---|---|---|---|
| 15 | E37K | Indirect disrupting | 8 | E37 participates in numerous contacts, including an intramolecular salt-bridge with ACE2 R393 and a sidechain-mainchain polar contact with K353 on the ACE2 b-hairpin that contributes to the binding interface. E37K may unfix the position of b-hairpin relative to a1 such that K353 no longer, participates in a polar contact with RBD G396. |
| 16 | D38V | Direct disrupting | 1 | Loss of two strong polar contacts with CoV2 RBD Y449. |
| 17 | Y50F | Direct disrupting | 2 | Loss of polar contact with RBD N487but may retain some hydrophobic packing with F486. |
| 18 | N51S | Indirect disrupting | 1 | Located on the C-terminal end of a1, N51 potentially helps anchor the helix through backbone-backbone polar contact with ACE2 L359 and backbone-backbone sidechain-backbone interactions with S47. Loss of these interactions could result destabilize a1 concavity which may alter affinity. |
| 19 | K68E | Indirect disrupting | 2 | Located on the a2 helix, K68 may be important for helix-helix packing. K68E could disrupt the packing critical for the binding interface. |
| 20 | F72V | Indirect disrupting | 1 | Located on the a2 helix, F72 participates in hydrophobic interactions and appears important in positioning of a1 helix. |
| 21 | Y83H | Direct disrupting | 2 | Loss of polar contact with RBD N487 |
| 22 | G326E | Direct disrupting | 1 | Induces steric clash with RBD G502 |
| 23 | G352V | Direct disrupting | 1 | Steric clash with RBD Y505 |
| 24 | D355N | Indirect disrupting | 2 | D355 is critical for the stability of the b-hairpin loop that forms part of the binding interface. |
| 25 | Q388L | Indirect disrupting | 4 | Q388 stabilizes a loop proximal to the ACE2-RBD interface through sidechain-backbone polar contact with ACE2 L560. Q388L would diminish this interaction. |
